# Chemo-phosphoproteomic profiling with ATR inhibitors berzosertib and gartisertib uncovers new biomarkers and DNA damage response regulators

**DOI:** 10.1101/2023.04.03.535285

**Authors:** Rathan Jadav, Florian Weiland, Sylvie M. Noordermeer, Thomas Carroll, Yuandi Gao, Jianming Wang, Houjiang Zhou, Frederic Lamoliatte, Ivan Muñoz, Rachel Toth, Thomas Macartney, Fiona Brown, C. James Hastie, Constance Alabert, Haico van Attikum, Frank Zenke, Jean-Yves Masson, John Rouse

## Abstract

The ATR kinase protects cells against DNA damage and replication stress and represents a promising anti-cancer drug target. The ATR inhibitors (ATRi) berzosertib and gartisertib are in clinical trials for treatment of advanced solid tumours as monotherapy or in combination with genotoxic agents. However, the pharmacodynamic ATR biomarker phospho-CHK1 has shown limited sensitivity in for quantitative assessment of ATR activity in clinical trials. Therefore, better biomarkers are needed, and with this in mind we carried out quantitative phospho-proteomic screening for ATR biomarkers that are highly sensitive to berzosertib and gartisertib. Screening identified novel ATR-dependent targets in three broad classes: i) targets whose phosphorylation is highly sensitive to ATRi; ii) novel targets with known genome maintenance roles; iii) novel targets whose cellular roles are unclear, including SCAF1. We show that SCAF1 interacts with RNAPII in a phospho-dependent manner and suppresses homologous recombination in cells lacking the *BRCA1* tumour suppressor. Taken together these data reveal potential new ATR biomarkers and new genome maintenance factors.

## Introduction

The ATR protein kinase plays critically important roles in the maintenance of genome stability (1–3). Through its targeting subunit ATRIP, ATR is recruited to sites in the genome where replisome progression is impeded (sites of replication stress; RS) (4–6). Stalled replication forks are vulnerable in nature, prone to degradation and collapse, and a host of proteins, including ATR, are dedicated to protecting these structures so that replication of the affected genomic region can continue once the replisome-blocking impediment has been removed or bypassed (1, 7). Full loss of ATR causes cell (and organism) lethality, probably because of the catastrophic chromosome shattering during S-phase (6, 8). Hypomorphic mutations in ATR cause diseases such as Seckel syndrome (9).

ATR belongs to the PI 3-kinase-related kinases (PIKKs) family which includes the double-strand-break (DSB)-activated kinases ATM and DNA-PK (10, 11). All three kinases are recruited to sites of DNA damage (ATM, DNA-PK, ATR) or fork stalling (ATR) by associated subunits, where they are activated enabling phosphorylation of target proteins on Ser/Thr-Gln (S/T-Q) motifs that together help to protect genome stability (3, 12–14). A major target of ATR is CHK1, which is itself a kinase activated by ATR-mediated phosphorylation on several residues including Ser345 (15, 16). Together ATR and CHK1 play key roles in the stabilization of replication forks, and activation of cell cycle checkpoints to prevent entry to mitosis in the presence of excess replication stress (17). In addition to its role as a key RS regulator, ATR is involved in inter-strand crosslink (ICL) repair, telomere control, DNA double strand break (DSB) repair and meiosis (18–21). A range of targets have been described for ATR beyond CHK1 and some of these targets are involved in ATR signalling including TOPBP1, RAD17, the RPA heterotrimer and ATRIP (10, 22).

Although it is an essential kinase, ATR has emerged over the years as a promising anti-cancer drug target (23–26). Seminal early studies identified small molecule ATR inhibitors (ATRi) such as ETP-46464 allowing a series of proof-of-concept studies which validated ATR as a drug target (27). These inhibitors, in conjunction with studies in ATR hypomorphic mice, helped to demonstrate that even though ATR activity is crucial in dealing with the low levels of RS in proliferating healthy cells, its activity becomes more important in tumour cells harbouring activated oncogenes such as cyclin E (*CCNE1*), MYC and RAS (27–29). This may reflect the elevated levels of RS in tumours resulting from the disruption of cell cycle regulation. In this light, several reports revealed that ATR inhibition is selectively toxic to tumours with high levels of DNA damage and RS (30–33). Moreover, ATR inhibition was shown to be toxic in cancer cells harbouring ATM mutations, a feature seen in many tumours (34–36), especially in combination with PARP inhibitors (35, 37). These observations stem from the loss of the p53-dependent DNA damage-induced G_1_ checkpoint in ATM-deficient cells, rendering them highly reliant on the S- and G_2_/M DNA damage checkpoint controlled by ATR. This raises the question - would ATR inhibition be toxic for healthy tissues? ATR hypomorphic mice expressing only low levels of ATR are viable and surprisingly normal (28, 29). Thus, it appears low levels of ATR activity are sufficient for cell health unless the levels of RS rise above a threshold, for example upon oncogene activation. These considerations have made ATR attractive as an anti-cancer drug target.

Berzosertib (formerly known VE-822, VX-970 and M6620) as was the first ATR inhibitor to be tested in humans for anti-cancer activity. Originally developed by Vertex as VE-822 and later as VX-970, it is a potent and highly selective inhibitor of ATR, with an IC_50_ of 0.2 nm *in vitro* and around 2 nM in cells (38). Several studies have shown that berzosertib potently sensitizes certain cancers to radio- and chemotherapy, with enhanced selectivity over normal cells (38–44). Berzosertib works particularly well in combination with the DNA crosslinking drug cisplatin, and in preclinical lung cancer xenograft models berzosertib induced tumour regression in combination with cisplatin (39, 44, 45). Given the potent anti-cancer activity in pre-clinical models, berzosertib has entered at least 19 clinical trials – phase I and II – since the first trial in 2012 (24). Most of these are designed to test berzosertib in combination with other drugs, but two of the trials will evaluate berzosertib as a monotherapy in a range of solid tumours, in particular those tumours with loss-of-function ATM mutations. Other ATR inhibitors have also entered clinical trials including the Astra Zeneca drug ceralasertib (AZD6738) and the Merck drug gartisertib (previously known as VX-803, M1774 and M4344) which is structurally unrelated to berzosertib. Gartisertib, the most potent ATR inhibitor known with an IC50 *in vitro* of 0.15 nM, was found to show synergy with DNA damaging chemotherapeutic agents and efficacy in xenograft models(46). It was entered into phase I clinical trials as monotherapy or in combination with carboplatin in patients with advanced solid tumours (NCT02278250).

Pharmacodynamic biomarkers are important for evaluating target engagement in clinical trials, and phospho-CHK1 (pSer345) has worked as a biomarker to monitor ATR inhibition after berzosertib administration in combination with cisplatin to treat advanced solid tumours including PARP inhibitor-resistant *BRCA1*-mutated germline ovarian cancer and metastatic colorectal cancers with *ATM* or *ARID1A* mutations (47). However, this pCHK1 biomarker was insufficiently sensitive to monitor ATR activity under basal conditions without genotoxic drug administration. Therefore, more sensitive biomarkers are needed. With this in mind, we employed a high-sensitivity, quantitative phospho-proteomic pipeline and carried out a global screen for proteins whose phosphorylation is inhibited by berzosertib or gartisertib, reasoning that hits in common are unlikely to be off–target. These analyses generated rich datasets, revealing a wide range of new ATR and CHK1 targets, and new players in the cellular response to DNA damage.

## Results

### Optimizing ATR activation and inhibition

We set out to establish optimal conditions for activating and inhibiting ATR in U-2 OS osteosarcoma cells. We first synchronized cells in S-phase, as ATR is activated at this phase of the cell cycle. To this end, cells were released from a 24h thymidine block into nocodazole for 12h; approximately 11h after release from the nocodazole-induced G_2_ arrest, the majority of cells were in S-phase (Fig. 1A). As shown in Fig. 1B, addition of hydroxyurea to S-phase synchronized cells caused a higher level of ATR activation compared with asynchronous cells, judged by CHK1 Ser345 phosphorylation (compare lane 2 with lane 4). To avoid inducing DSB (and activating ATM and DNA-PK indirectly) we sought to use the lowest HU exposure time necessary to fully activate ATR, and the lowest dose of ATRi needed for full inhibition. As shown in Fig. 1C, a 30 min HU treatment was sufficient to activate ATR, judged by CHK1 (pSer345), and pre-incubation of cells with 1μM berzosertib or gartisertib for 1 hr prior to HU was sufficient to fully block phosphorylation of CHK1 pSer345 (Fig. 1B). Under these conditions, no increase in phosphorylation of H2AX, a marker associated with DSB formation, or phospho-RPA, a marker of DNA end resection, was detected (Fig. 1C). Based on these data, we settled on the cell treatment workflow shown in Fig. 1D for phosphoproteomic screening.

**Figure 1.**
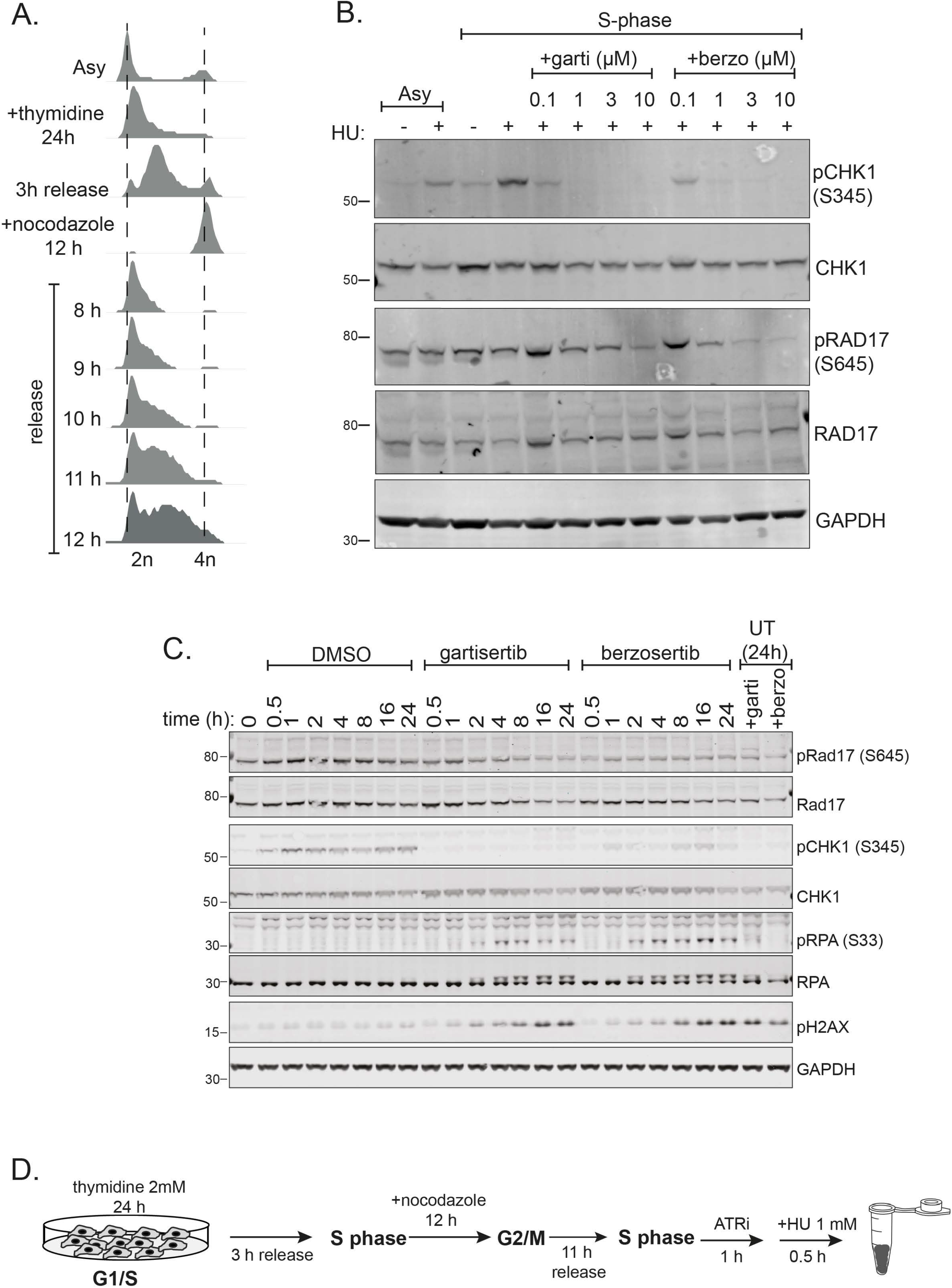
Optimizing activation and inhibition of ATR. A. U-2 OS cells were incubated with thymidine (2 mM) for 24h and released for 3h at which point nocodazole (100 ng/ml) was added for a further 12h. Cells were released from nocodazole into fresh medium for the times indicated. Cells were fixed, stained with propidium iodide (PI) and analysed by FACS. B. U-2 OS cells synchronized in S-phase (11h after release from nocodazole) were pre-incubated for 1h with the indicated concentrations of berzosertib or gartisertib before addition of HU (1 mM) for 1h. Cells were lysed, and extracts subjected to western blotting with the antibodies indicated. C. Same as B., except that S-phase cells were pre-incubated with berzosertib or gartisertib (1 μM for 1h) before addition of HU (0.5 mM) for the times indicated. D. Optimized cell treatment workflow.

### Phospho-proteomic screening for phosphorylation events inhibited by berzosertib and gartisertib

We first carried out a quantitative phospho-proteomic screen with cells exposed to [HU+DMSO] or [HU+berzosertib] according to the pipeline shown in Fig. S1A. Five biological replicates of each of the two cell populations were lysed (Fig. S1B, left panels), and Cys residues were reduced and alkylated. After trypsinization of cell extracts, phosphopeptides were enriched by chromatography. The ten samples were then isotopically labelled with tandem mass tags (TMT), allowing multiplexed and quantitative analysis of all ten samples which were combined and analysed in parallel (48). Applying a false discovery rate (FDR) threshold of 5% identified 21,178 unique phospho-peptides of which 17,128 had at least one phosphorylation site with a localisation probability of ≤ 75% (14,251 unique sites); this yielded 9,367 unique phosphorylation sites with a 1% false localization rate (Tables S1, S2) as called by MaxQuant. Normalisation and intensity distribution in the TMT channels was checked and deemed satisfactory (Figs. S1C, E). To define berzosertib-sensitive phosphorylation events, mass spectrometric data were visualized in a volcano plot, which revealed 553 phosphopeptides (463 unique sequences) that were lower in abundance after exposure of cells to berzosertib (Fig. 2A, Table S2); all the phosphopeptides within this group had an adjusted p-value of < 0.05 (5% FDR).

**Figure 2.**
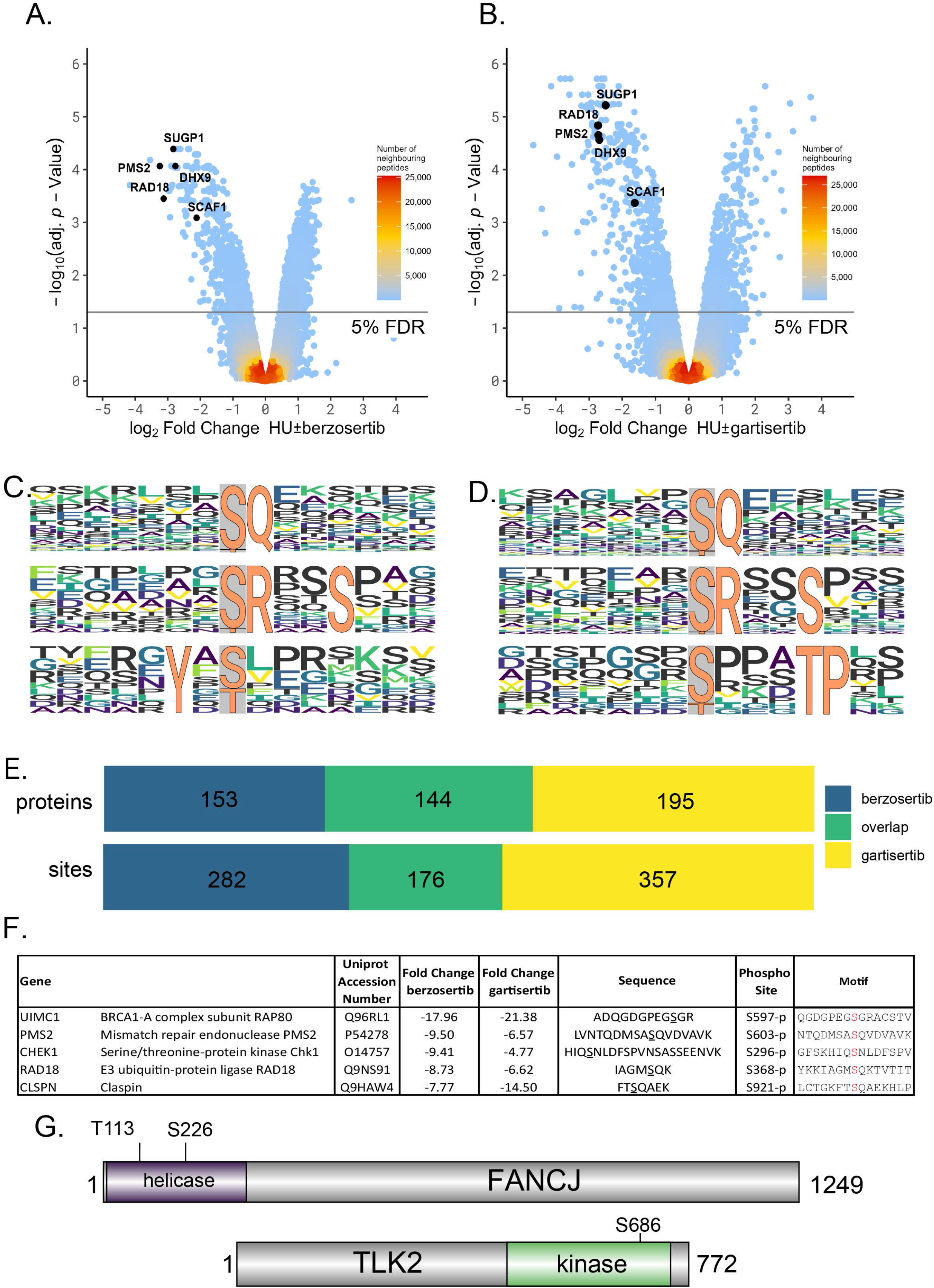
Phosphoproteomic screening of phosphorylation sites sensitive to berzosertib or gartisertib. A, B. Volcano plot showing phosphorylation sites affected by berzosertib (A) or gartisertib (B). The horizontal cut-off lines represent an adjusted p-value of 0.05.Phospho-peptides lower abundant under inhibitor treatment are in the negative logFC region of the plots. The mass spectrometry proteomics raw data for this figure have been deposited to the ProteomeXchange Consortium (109) via the jPOSTrepo partner repository (108) with the dataset identifier PXD040469. Data analysis scripts are available via Zenodo under https://doi.org/10.5281/zenodo.7784890. C, D. Phosphomotif analysis for berzosertib (C) and gartisertib (D). E. Overlap of proteins and phosphorylation sites affected by berzosertib or gartisertib. (F) The five proteins whose phosphorylation is most strongly affected by berzosertib (“Class 1” hits). The phosphorylated residue is highlighted in red in the “Motif” column. Fold change refers to the difference between HU±ATRi. G. Schematic diagram showing the location of the ATR-dependent phosphorylation sites in the DNA helicase FANCJ and the protein kinase TLK2.

A similar phosphoproteomic screen was carried out with cells exposed to [HU+DMSO] or [HU+gartisertib] according to the pipeline shown in Fig. S1A, again with five biological replicates per condition (Fig. S1B, right panels). Applying a FDR of 5% identified 22,853 unique peptides of which 17,743 had at least one phosphorylation site with a localisation probability of ≤ 75% (14,456 unique sites); this yielded 8,924 unique phosphorylation sites with a 1% false localization rate (Tables S1, S3). Normalisation and intensity distribution in the TMT channels was checked and deemed satisfactory (Figs. S1D, F). To define gartisertib-sensitive phosphorylation events, mass spectrometric data were visualized in a volcano plot, which revealed 657 phosphopeptides (559 unique sequences) that were lower in abundance after exposure of cells to gartisertib (Fig. 2B; Table S3); all the phosphopeptides within this cluster had an adjusted p-value of < 0.05 (5% FDR).

### Enrichment of a novel phospho-motif

Analysis of the amino acid sequences surrounding the phosphorylation sites inhibited by berzosertib and gartisertib revealed strong enrichment of two different phospho-motifs common to both inhibitors: the pS/pT-Q motif typical of ATR and other PI-(3) kinase-like kinases, and a p<u>S</u>RXXS motif where the first serine is the phosphorylated residue (Figs. 2C, D). This latter motif is presumably targeted by an ATR-activated kinase other than CHK1 which phosphorylates S/T residues in an RXX<u>S</u> motif *in vitro* and in cell extracts; it should not phosphorylate the first Ser in the SRXXS motif (49–51). Intriguingly, each ATRi inhibited phosphorylation of Ser/Thr residues in a third phospho-motif that was specific to each inhibitor: YX[pS/pT] for berzosertib and pSXXXTP for gartisertib (Figs. 2C, D). It is possible these motifs reflect off-target effects uniquely associated with each inhibitor. Gene ontology (GO) analysis showed a striking enrichment of the terms DNA repair, DNA replication, double-strand break repair (NHEJ and HR) for both ATRi (Fig. S2A). Comparison of the phosphorylation sites inhibited by berzosertib or gartisertib revealed an overlap of around 45% corresponding to 176 sites in 144 target proteins (Fig. 2E). The top 40 hits common to both ATRi are listed in Table 1, the full list is given in Table S4. Given these sites are common to the two different ATRi we used, we regard them as *bona fide* ATR-dependent phosphorylation events.

**Table 1.**
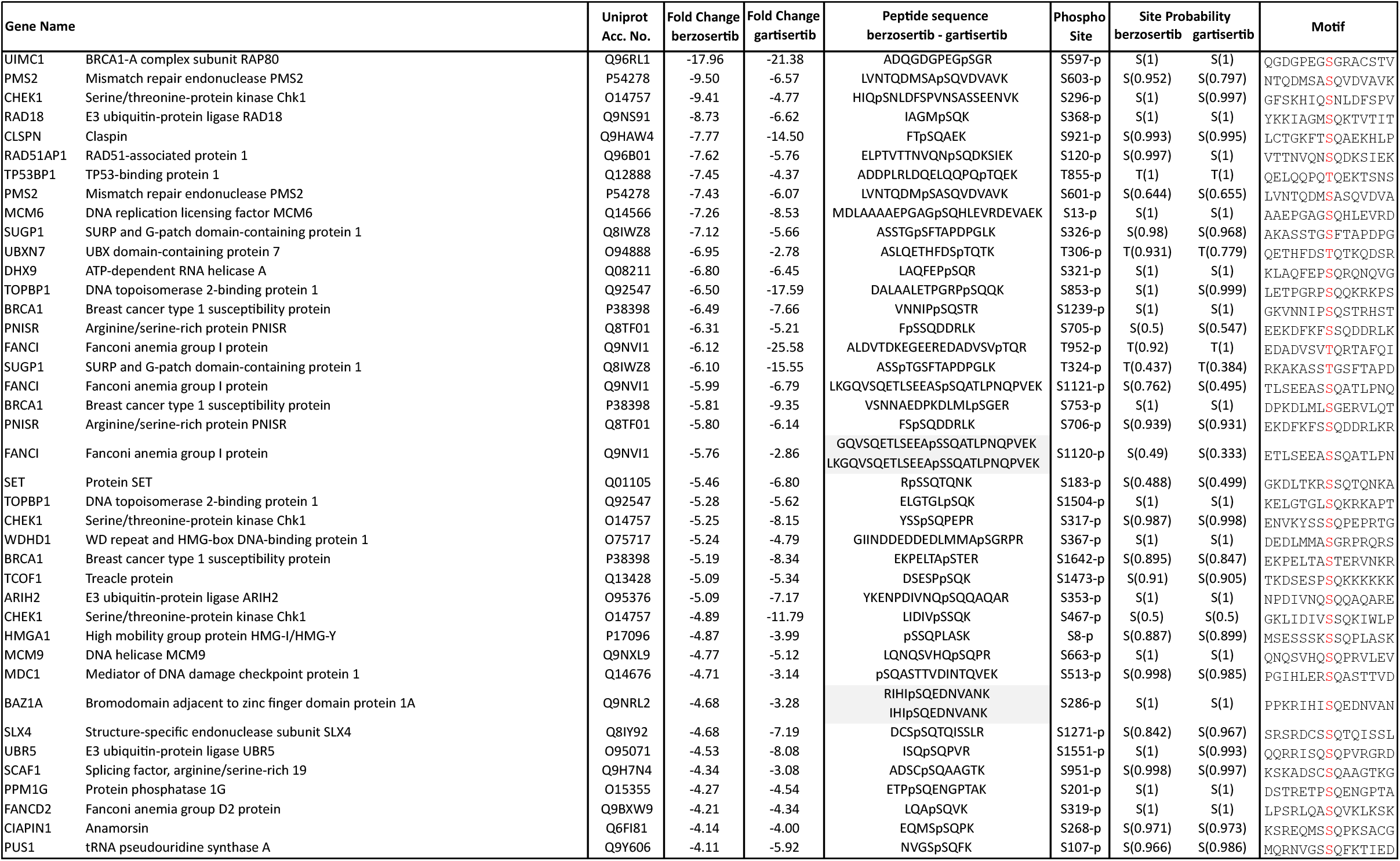
ATRi phosphoproteomic dataset: top 40 hits. List of the top 40 phosphorylation sites inhibited by both berzosertib and gartisertib, ranked according to fold-change with berzosertib. Phosphorylation sites are highlighted in red in the “Motif” column.

### ATRi-sensitive targets fall into three classes

We divided the phosphorylation sites sensitive to both ATRi into three classes. The first class are those showing the highest degree of sensitivity to ATRi, and these sites have good potential as new ATR biomarkers. The top 5 berzosertib-sensitive sites, that are also sensitive to gartisertib, are shown in Fig. 2F. The most sensitive phospho-site is pSer597 of RAP80, a subunit of the BRCA1-A complex (52), which displayed a fold change in phosphorylation of 21 and 18 with berzosertib and gartisertib respectively. This was not previously known as an ATR-dependent phospho-site. PMS2 pSer603 and RAD18 pSer368, which are also among the top 5 phosphorylation sites with the highest sensitivity to ATRi, both lie in classical SQ motifs and are therefore likely to be direct ATR targets. Also featuring in the top 5 was CHK1 pSer296, a known DNA damage-induced autophosphorylation site (53).

The second class of ATR-dependent phosphorylation sites are those found in proteins already implicated in cellular DNA damage responses, but that were not known to be ATR targets (Table S5). Examples of proteins in this category include: MCM9 which interacts with MCM8 to form a heterohexamer (paralogous to the MCM2-7 replicative helicase) involved in post-synaptic DNA synthesis during homologous recombination (HR) (54); the SLX4 scaffold protein which tethers and coordinates three structure-selective DNA repair nucleases (SLX1, XPF–ERCC and MUS81–EME1) (55); RAP1 (TERF2IP), which interacts with the shelterin component TRF2 to regulate telomere length and protection (56). Of particular interest in this category are novel ATR-catalysed phosphorylation sites within functionally annotated catalytic domains. For example, FANCJ is a BRCA1-associated, DNA-dependent ATPase and helicase involved in HR and ICL repair. (Fig. 2G; Table S5) (57, 58). In this study we found that FANCJ is phosphorylated on two residues in the helicase catalytic domain – Thr113 and Ser226, both of which conform to the classical S/T-Q ATR consensus motif. The kinase TLK2, implicated in chromatin assembly, replication fork integrity and recovery from DNA damage induced G_2_ arrest is phosphorylated on Ser686 (Fig. 2G; Table S5) (59, 60). This is a highly conserved residue within the kinase catalytic domain, raising the possibility that ATR regulates TLK2 kinase activity.

We also noticed that the DHX9 helicase is phosphorylated in an ATR-dependent manner on Ser321 which lies in a classical SQ consensus motif (Tables 1 & S5). DHX9 is a poorly understood helicase capable of unwinding DNA, RNA and hybrid nucleic acids *in vitro*, with pleiotropic roles in maintenance of genome stability (61). Recently, DHX9 was shown to facilitate R-loop formation and to stimulate BRCA1-dependent DNA end resection and HR (62, 63). Ser321 lies close to the start of the helicase domain suggesting it may influence DHX9 activity and/or function (Fig. 3A), and we next sought to validate ATR-dependent phosphorylation of this site on DHX9 expressed in U-2 OS cells. Extracted ion chromatogram (XIC) analysis of tryptic phosphopeptides isolated from GFP-tagged DHX9 (pSer321) confirmed that the HU-induced phosphorylation of this site is reduced by preincubating cells with berzosertib but not with the CHK1 inhibitor PF477736 (64) (Figs. 3B, C). Therefore, DHX9 is a target of ATR.

**Figure 3.**
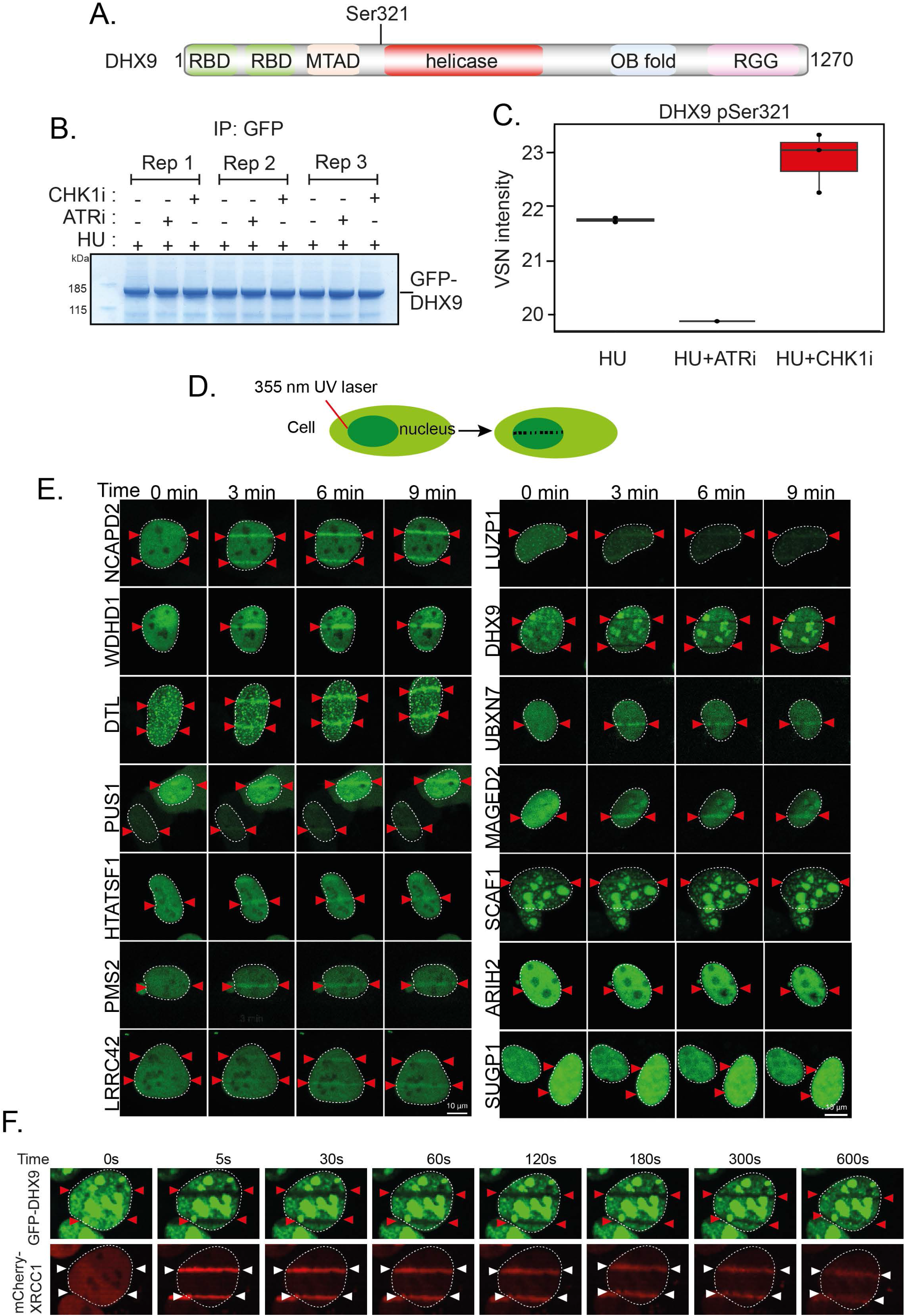
Screening novel ATR targets for recruitment to DNA damage sites. A. Schematic diagram showing the domain organization of DHX9. RBD, RNA binding domain; MTAD, minimal transcriptional activation domain; RGG, RGG-rich domain; OB fold, oligonucleotide/oligosaccharide binding fold. B. U2 O-S cells were transfected with GFP-tagged DHX9 and after 24h cells were lysed, and cell extracts were subjected to immunoprecipitation with anti-GFP-agarose beads. Precipitates were subjected to SDS-PAGE and staining with Coomassie Brilliant Blue; the bands corresponding to the GFP-DHX9 were excised and processed for mass spectrometric detection of relevant phospho-peptides. Three independent co-transfection experiments were done for every condition (Rep=biological <u>rep</u>licate). C. Label free quantification was used to generate a boxplot showing VSN transformed intensity of phospho-peptides containing to DHX9 pSer^321^. Mass spectrometry raw data was uploaded to ProteomeXchange via the PRIDE partner repository (111) and can be downloaded via the identifier PXD041250. Data analysis scripts can be accessed via Zenodo under https://doi.org/10.5281/zenodo.7788447. D. Schematic diagram showing micro-irradiation of BrdU-sensitized cells to induce DNA damage along a track in the nucleus. E. BrdU–sensitized U–2–OS cells transiently expressing GFP-tagged forms of the proteins indicated were line micro-irradiated and imaged after 2 min. F. BrdU-sensitized U-2 OS cells stably expressing mCherry XRCC1 and expressing GFP-DHX9 in a tetracycline inducible manner were subjected to laser micro-irradiation and live imaged at the times indicated.

The third class of ATR-dependent phosphorylation sites are found in proteins that were not previously linked to cellular responses to DNA damage or RS (Table S6). Several proteins are of unknown function, and we focussed on some of these.

### Secondary screening: recruitment to DNA damage sites

All of the proteins in the third class of ATR targets mentioned above are potentially new DDR proteins. Re-localization to DNA damage sites is a universal feature of proteins involved in DDR, and we next tested a range of class 3 proteins for this behaviour. Cells pre-labelled with BrdU expressing GFP-tagged versions of each protein were subjected to micro-irradiation to induce DNA damage along a track in the nucleus with a 355 nm laser (Fig. 3D). Some of the proteins tested showed robust recruitment to DNA damage sites (Figs. 3E and S2B). For example, the condensin subunit NCAPD2, the replisome subunit WDHD1/AND1, pseudouridine synthase PUS1, and the E3 ubiquitin ligase ARIH2 are recruited rapidly and in a sustained manner to DNA damage sites and may therefore play previously unanticipated roles in the DDR (Figs. 3E and S2B). DHX9 was the only protein we found to be excluded from DNA damage sites (Fig. 3F); the underlying mechanism is not yet known but exclusion is not prevented by berzosertib or gartisertib (data not shown).

Several ATR targets of unknown function are recruited to micro-irradiation tracks, strongly suggesting roles in DDR. Some of these showed similarity in the kinetics of recruitment – for example SUGP1 and LUZP1. SUGP1 is an uncharacterized protein containing two SURP motifs often found in proteins involved in pre-mRNA splicing (65–67), and a G-patch motif found in RNA binding proteins and in particular those with SURP motifs (68) (Fig. 4A). LUZP1 is largely uncharacterized but has been implicated recently in control of primary cilia (Fig. 4A) (69–71). Recruitment of GFP-tagged forms of both of these proteins to micro-irradiation sites was rapid and transient (Figs. 4B, C), reminiscent of proteins that bind poly–ADP ribose (PAR) chains generated by DNA damage–activated poly–ADP ribose polymerases (PARPs) (72). Consistent with this idea, recruitment of LUZP1 and SUGP1 was blocked by the PARP inhibitor olaparib; in contrast, retention time was prolonged by PDD00017273, an inhibitor of PARG (poly–ADP ribose glycohydrolase) which delays PAR degradation (Figs. 4B, C) (73).

**Figure 4.**
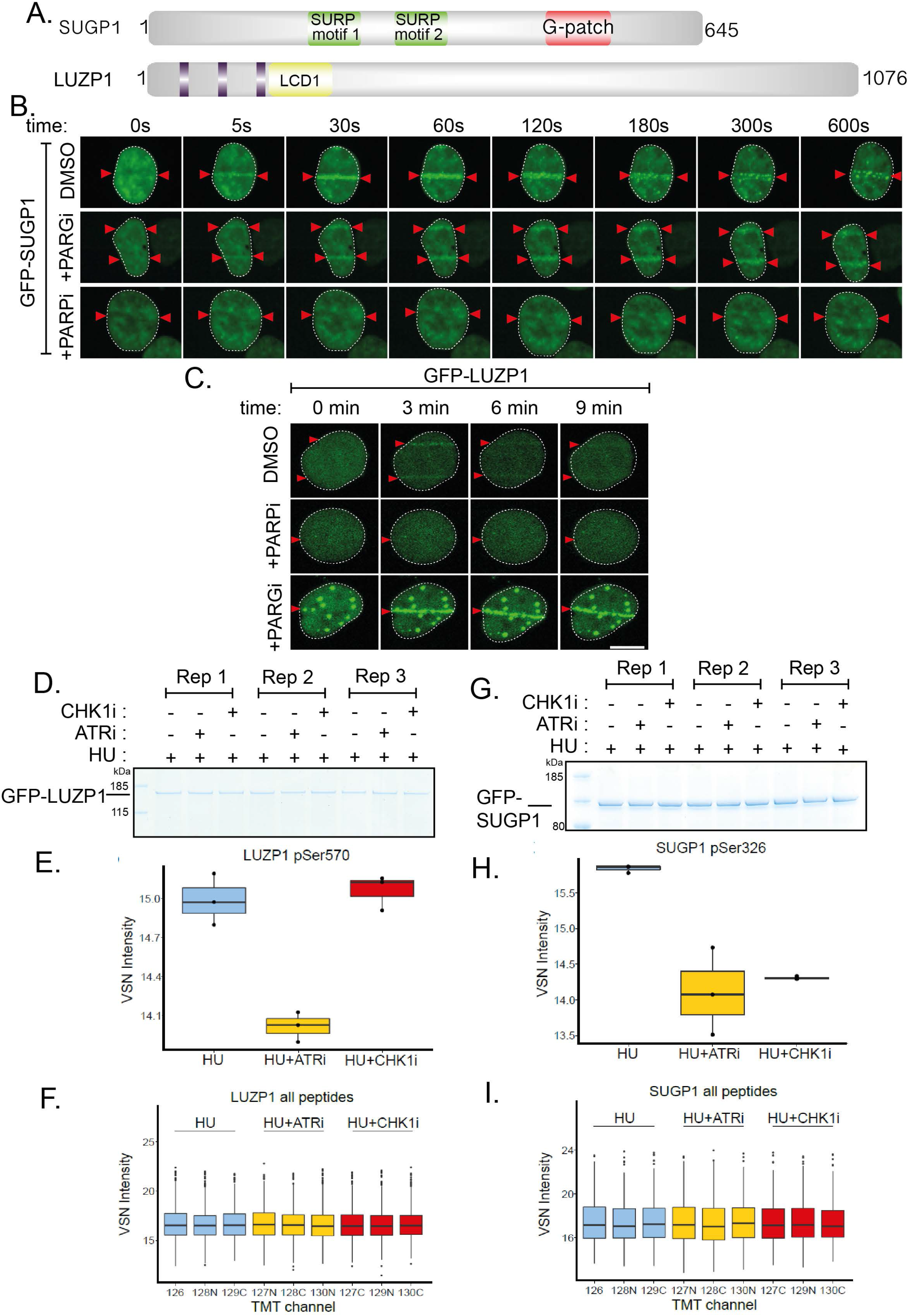
Phosphorylation of SUGP1 and LUZP1 and recruitment to DNA damage sites. A. Schematic diagram showing the domain organization of SUGP1 and LUZP1. B, C. U-2 OS cells stably expressing GFP–SUGP1 (B) or GFP–LUZP1 (C) were pre– incubated with DMSO (mock), olaparib (5 μM; PARPi) or PDD00017273 (0.3 μM; PARGi) for 1 h prior to line micro–irradiation. Cells were live imaged at the times indicated. D. U2 O-S cells were co-transfected with GFP-LUZP1. After 24h cells were lysed, and cell extracts were subjected to immunoprecipitation with anti-GFP-agarose beads. Precipitates were subjected to SDS-PAGE and staining with Coomassie Brilliant Blue, and the bands corresponding to the GFP-tagged proteins were excised and processed for mass spectrometric detection of relevant phospho-peptides. Three independent co-transfection experiments were done for every condition (Rep=biological <u>rep</u>licate). E. Boxplots showing VSN–normalised intensity of phospho-peptides corresponding to LUZP1 pSer570 from the experiments in (D). F. Boxplots of the VSN-adjusted TMT reporter ion intensities for all peptides for each TMT label in the case of GFP–LUZP1 from the experiment in (D). G-I. Same as D-F except cells were transfected with GFP-SUGP1 and phosphorylation of Ser326 was analysed in a similar manner. Mass spectrometry raw data was uploaded to ProteomeXchange via jPOSTrepo and can be accessed under the identifier PXD040476. Accompanying data analysis scripts are available at Zenodo under https://doi.org/10.5281/zenodo.7661023.

We next sought to validate ATR-dependent phosphorylation of these proteins by testing the phosphorylation of these proteins expressed in U-2 OS cells. LUZP1 Ser326 lies in the new consensus phosphomotif enriched among phosphorylation sites inhibited by berzosertib and gartisertib: p<u>S</u>RXXS (Fig. 2C, D). Extracted ion chromatogram (XIC) analysis of tryptic phosphopeptides isolated from GFP-tagged LUZP1 (pSer570) confirmed that the HU-induced phosphorylation of this site is reduced by preincubating cells with berzosertib but not with the CHK1 inhibitor PF477736 (64) (Figs. 4D-F). These data imply LUZP1 Ser570 is phosphorylated by a kinase downstream of ATR that is not CHK1 (Table 1). Similar XIC analysis of SUGP1 (pSer326) revealed that HU-induced phosphorylation of this site is reduced by preincubating cells with berzosertib or PF477736 (Figs. 4G-I). Therefore, SUGP1 lies downstream of both ATR and CHK1 but pSer570 does not conform to the putative CHK1 RXXp<u>S</u> consensus motif, suggesting that a CHK1-activated kinase is responsible.

### Phospho-dependent interaction of SCAF1 with the RNAPII CTD

SCAF1 is a protein of unknown function that we decided to investigate further. Like SUGP1 and LUZP1, we found SCAF1 is recruited to sites of laser micro-irradiation in a transient manner; recruitment was blocked by olaparib, and retention time was prolonged by PDD00017273, demonstrating dependence of poly-ADP ribose formation (Figs. 5A, B). The cellular roles of SCAF1 are unknown, but it has a Set-Rpb1 interaction (SRI) domain towards the C-terminus (Fig. 5C) (74) which suggested a link to RNA polymerase (RNAP) II. The largest subunit of human RNAPII (POLR2A) has a C-terminal domain (CTD) bearing 52 tandem repeats of the consensus heptapeptide sequence Tyr^1^-Ser^2^-Pro^3^-Thr^4^-Ser^5^-Pro^6^-Ser^7^. CDK9/cyclin T preferentially phosphorylates the Ser2 sites of this sequence to promote transcriptional elongation; CDK7/cyclin H, a component of the general transcription factor (TFIIH), preferentially phosphorylates Ser5, which facilitates promoter clearance and transcriptional initiation (75, 76). SRI domains, in proteins such as the histone methyltransferase SETD2 and the helicase RECQL5 (implicated in DNA replication, transcription and repair), interact with the Ser2/Ser5-phosphorylated form of RNAPII CTD (74, 77–80). To test if the SCAF1 interacts with RNAPII, endogenous SCAF1 was immunoprecipitated from RPE-1 cells using antibodies generated in-house, and *SCAF1* knockout (KO) RPE-1 cells were used as control. Quantitative mass spectrometric analysis identified a range of proteins that were much higher in abundance in SCAF1 precipitates from parental cells than from SCAF-KO cells (Fig. 5D). Besides SCAF1 itself, this included POLR2A and several other subunits of RNAPII as well as RECQL5, known to associate with RNAPII through its SRI domain (79). We speculate the interaction of RECQL5 is indirect through RNAPII-CTD.

**Figure 5.**
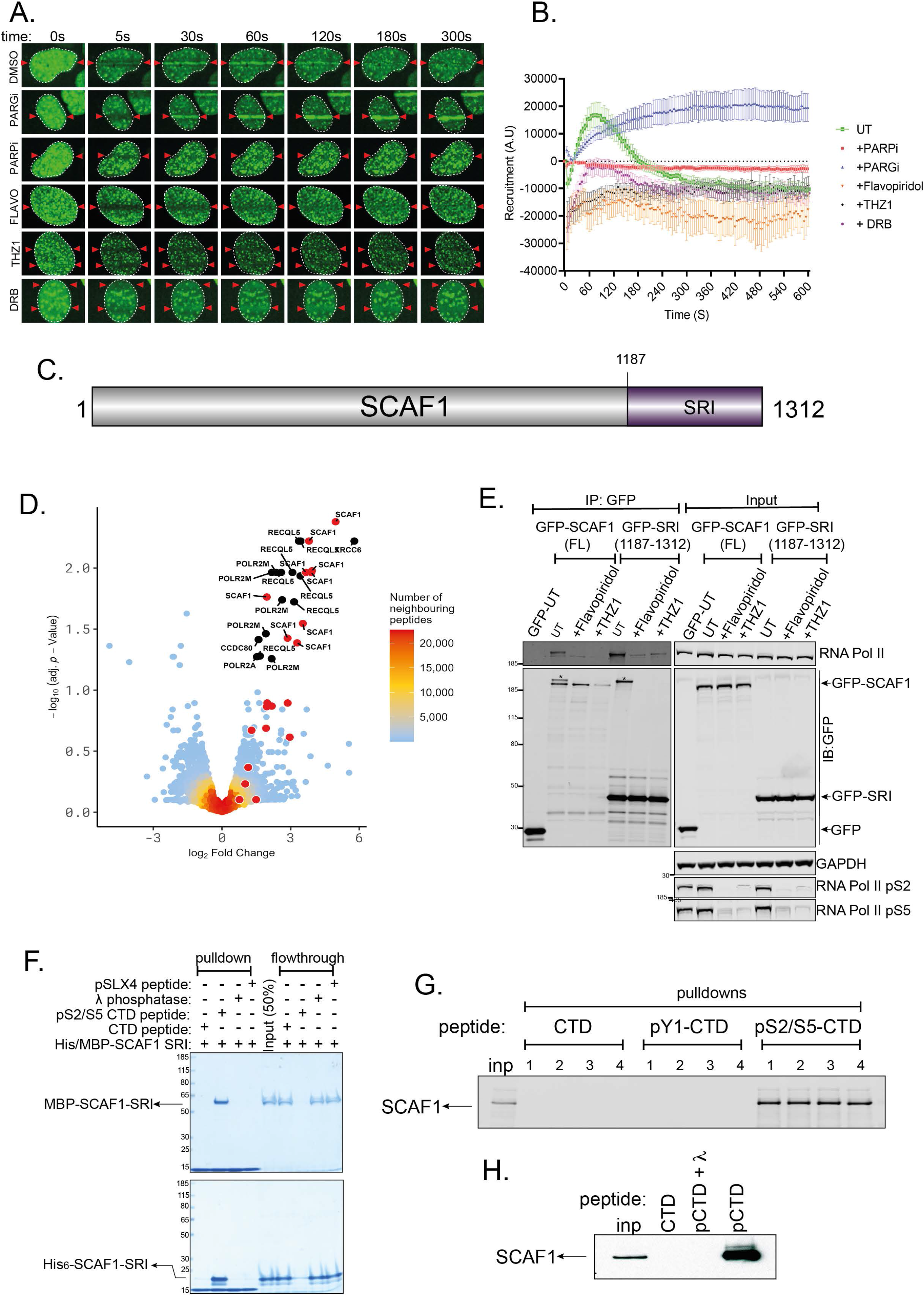
SCAF1 interacts with the Ser2/Ser5-phosphorylated CTD of RNAPII. A. BrdU–sensitized U2 O-S Flp–In T–REx cells stably expressing GFP-tagged SCAF1, pre–incubated with DMSO, olaparib (5 μM; PARPi), PDD00017273 (0.3 μM; PARGi), flavopiridol (10 μM), THZ-1 (10 μM) or DRB for 1 h were micro-irradiated with a 405 nm laser and imaged at the times indicated. B. Same as G. except that cells stably expressing GFP–SCAF1 subjected to spot micro–irradiation (405 nm), and spot intensities were quantitated. Data represent the mean ± SEM of two independent experiments; > 50 micro–irradiated cells per point. C. Schematic diagram showing the domain organization of SCAF1. D. Lysates of RPE1 *hTERT TP53^−/−^*cells parental cells and RPE1 *hTERT TP53^−/−^ SCAF1^−/−^*cells were subjected to immunoprecipitation with in-house sheep anti-SCAF1 antibodies (3 biological replicates). Proteins were eluted from beads, trypsinized, and after TMT labelling, samples were pooled and injected on an UltiMate 3000 RSLCnano System coupled to an Orbitrap Fusion Lumos Tribrid Mass Spectrometer. A volcano plot representing SCAF1 interactors is shown. Mass spectrometry raw data were uploaded to ProteomeXchange via jPOSTrepo with the identifier PXD041201. Data analysis scripts are available from Zenodo under https://doi.org/10.5281/zenodo.7784982. E. U-2 OS Flp-In TRex cells expressing GFP-SCAF1, or GFP-SCAF1-SRI domain were subjected to immunoprecipitation with anti-GFP antibodies and precipitates were subjected to western blotting with the antibodies indicated (“IP-GFP”). Input cell extracts were analysed in parallel. F. A biotinylated peptide containing three heptad repeats from the POL2RA CTD (CTD peptide), or the corresponding peptide that was phosphorylated at Ser2 and Ser5 in each heptad (pCTD) incubated or not with lambda phosphatase, was immobilized on streptavidin beads. A peptide corresponding to phosphor-Ser1238 from SLX4 was used as control. Beads were incubated with MBP-SCAF1-SRI or His6-tagged SRI expressed in bacteria. Beads were washed and subjected to SDS-PAGE and Coomassie staining. Both bead and supernatant “flowthrough” fractions are shown. G. The immobilized CTD and pS2/S5-CTD peptides were incubated with extracts of U-2 OS cells; after washing, precipitates were subjected to western blotting with SCAF1 antibodies. A CTD peptide where Tyr1 in each heptad was phosphorylated (pY1-CTD) was used as control. H. Same as E, expect that a the pS2/S5-CTD peptides was pre-treated or not with lambda phosphatase.

We next set out to test if the interaction of SCAF1 with RNAPII is mediated by phospho-dependent interaction of the SCAF1 SRI domain with the Ser2/Ser5-phosphorylated form of POLR2A CTD. As shown in Fig. 5E, brief exposure of cells to flavopiridol or THZ1 which inhibit CDK7 and CDK9-dependent phosphorylation of POLR2A Ser2/Ser5 (81–83) severely reduces association of GFP-SCAF1 or GFP-SRI with POLR2A. To investigate further, synthetic peptides containing three heptad repeats from the POL2RA CTD, phosphorylated (“pCTD peptide”) at Ser2 and Ser5 of each heptad, bearing biotin at the N-terminus, were immobilized on streptavidin beads; the non-phosphorylated peptide was used as control (“CTD peptide”). Pulldown experiments revealed that recombinant MBP-tagged or His_6_-tagged forms of the SCAF1 SRI domain expressed in bacteria were efficiently retrieved by the pCTD beads but not by the CTD beads or beads bearing a control SLX4 phosphopeptide (Fig. 5F). Furthermore, lambda phosphatase pre-treatment of the pCTD beads prevented retrieval of the SCAF1 SRI domain. We also found that the pSer2/Ser5-CTD beads, but not beads bearing the CTD peptide or the CTD peptide where Tyr1 in each of the three heptads was phosphorylated, could retrieve endogenous SCAF1 from cell extracts (Fig. 5G). Pre-treatment of the pSer2/Ser5 CTD beads with lambda phosphatase prevented retrieval of SCAF1 from cell extract (Fig. 5H). We also tested if the inhibitors that block its interaction with RNAPII CTD (Fig. 5E) also affect recruitment of SCAF1 to DNA damage sites. As shown in Figs. 5A and B, the recruitment of SCAF1 to sites of laser micro-irradiation is inhibited by THZ-1 and flavopiridol, and in fact SCAF1 appears to be excluded from DNA damage tracks when cells are exposed to these drugs. SCAF1 recruitment is also inhibited by DRB (5,6–dichloro–1–b-D-ribofuranosylbenzimidazole) which blocks RNAP II elongation (Figs. 5A, B). In contrast, flavopiridol, THZ-1 and DRB do not affect recruitment of the PAR-responsive protein XRCC1 (Fig. S3). Taken together the data above show that the SRI domain of SCAF1 interacts with the Ser2/Ser5-phosphorylated form of RNAPII, and this interaction facilitates SCAF1 recruitment to DNA damage sites.

### SCAF1 deficiency in *BRCA1*^−/−^ cells causes partial restoration of HR

SCAF1 was one of a range of genes identified previously in a genome-wide CRISPR-based screen for gene deletions that reverse the sensitivity of RPE-1 hTERT *BRCA1*^−/−^ *TP53*^−/−^ (*BRCA1*-KO cells) to PARP inhibitors (84). We first set out to validate these data. As shown in Fig. 6A, two different small guide (sg) RNAs targeting SCAF1 caused a modest but reproducible increase in the resistance of *BRCA1*-KO cells, but not the parental *TP53***^−/−^** cells, to the PARP inhibitor olaparib. A SCAF1-specific siRNA had a similar effect (Fig. 6B; Fig. S4A). The reversal of olaparib sensitivity suggests that depleting SCAF1 may restore HR in *BRCA1*-KO cells, and we set out to investigate this possibility by visualizing the recruitment of the RAD51 recombinase to IR-induced DSB, known be defective in *BRCA1*-KO cells (85). We found that siRNA-mediated depletion of SCAF1 using a series of individual siRNAs increased the proportion of *BRCA1*-KO cells with RAD51 foci with similar effect size to depletion of 53BP1 (Figs. 6C, D; Figs. S4B, C), which is a well-known inhibitor of HR in *BRCA1*-deficient cells (86, 87). We used genome editing to generate clonal *SCAF1* gene knockouts in *BRCA1*-KO cells (Fig. 6E) and found that *SCAF1* deletion increased the proportion of cells with greater than five IR-induced RAD51 foci (Fig. 6E). Together these data indicate that SCAF1 suppresses HR in *BRCA1*-deficient cells.

**Figure 6.**
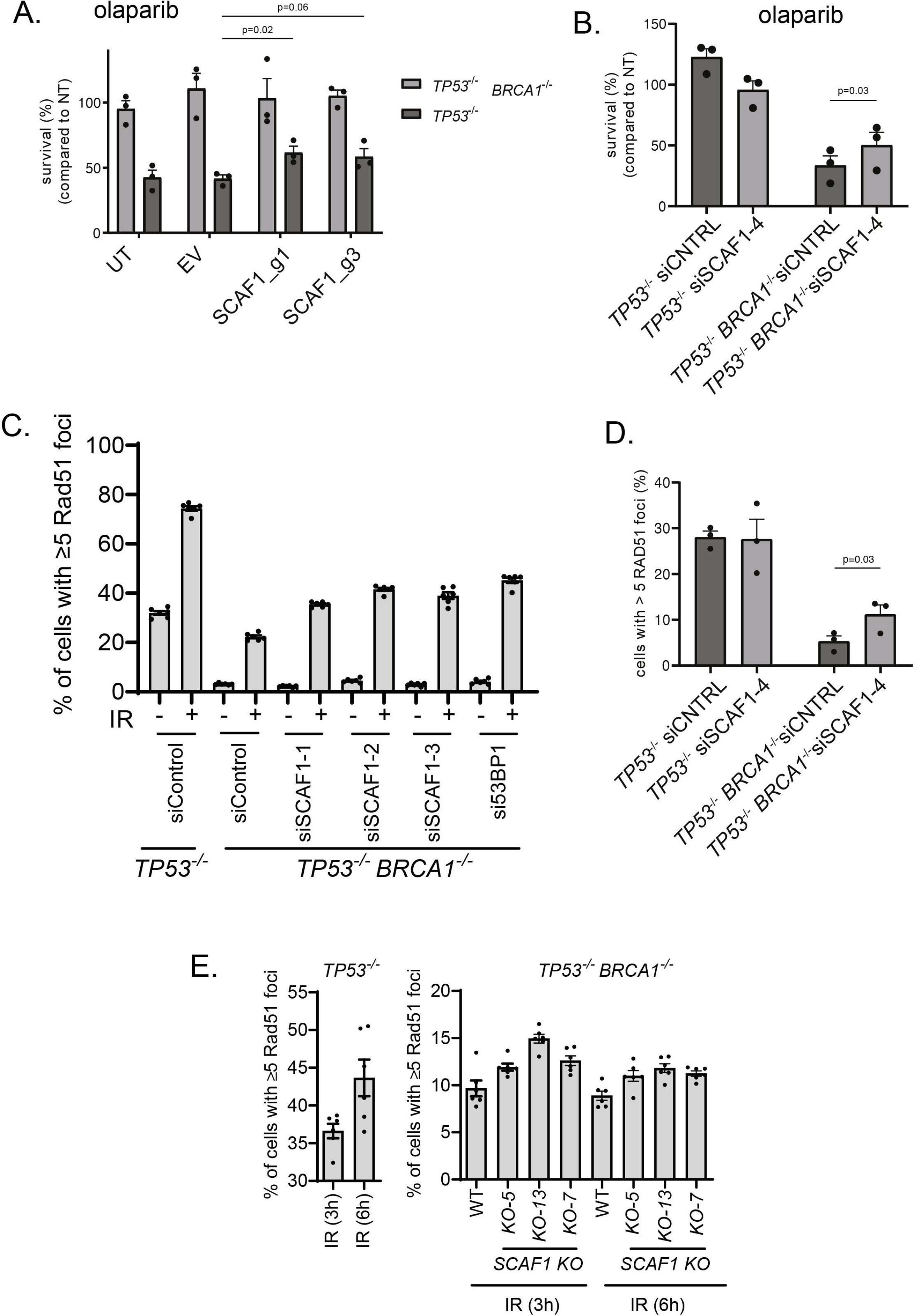
SCAF1 is a new genome maintenance factor. A. The cell lines indicated were virally transduced with sgRNAs targeting *SCAF1* and clonogenic survival was assessed in presence or absence of olaparib (16 nM) on the pool of transduced cells. sgRNA targeting efficiency was assessed using genomic PCR amplification of the targeted locus and TIDE analysis. Editing efficiencies were > 50% for all conditions (see Materials and Methods). Data is represented as mean + SEM (n=3), p-values are obtained using a two-tailed t-test. UT, untreated; EV, empty vector. B. Same as (A), but here SCAF1 was depleted using siRNA transfection prior to clonogenic survival. Data is represented as mean + SEM (n=3), p-value is obtained using a two-tailed t-test. C. The cell lines indicated were transfected with the siRNAs indicated and after 24h cells were exposed to IR (10 Gy). Cells were allowed to recover for 3h, fixed and the proportion of RAD51 cells with greater than five foci was assessed by immunofluorescence. Data was acquired with a high-content screening station ScanR. Data from three independent experiments was combined; data are represented as mean ± SEM. D. Same as C. expect that the experiment was performed by SN. Data is represented as mean + SEM (n=3), p-value is obtained using a two-tailed t-test. E. Same as C. except that the indicated cell lines were used.

The early steps of homologous recombination involve the resection of DNA ends (5′– 3′ degradation) leading to a stretch of 3’-ended single-stranded DNA which is then repaired using the sister chromatid as a DNA template. It was shown previously 53BP1 and shieldin suppress HR in *BRCA1*-KO cells by suppressing DNA end resection and we therefore analyzed the impact of SCAF1 on DNA resection. Resected DNA exposes BrdU-incorporated regions of single-stranded DNA, and immunofluorescence analysis using BrdU antibodies under non-denaturing conditions serves as a readout for DSB resection. We observed that deletion of *SCAF1* in *BRCA1*-KO cells leads to increased levels of single-stranded DNA (Fig. S4D, E). To further validate these results, we monitored phosphorylation of RPA32 at Ser 4/Ser 8, a surrogate marker of DNA resection. Surprisingly, the levels of RPA32 phosphorylation in *BRCA1*-KO *SCAF1*-KO cells were not noticeably different from single *BRCA1*-KO cells monitored by western blotting or immunofluorescence (Figs. S4F, G). These data suggest that, while increasing single-stranded DNA in BRCA1-KO cells, other mechanisms rather than DNA end resection may provide PARPi resistance after *SCAF1* deletion.

## Discussion

In this study we carried out parallel phosphoproteomic screens to identify proteins that are sensitive to the ATR inhibitors berzosertib or gartisertib. We identified 176 phosphorylation sites in 144 proteins whose phosphorylation is reduced by both drug, which corresponds to around 45% of the total number of phospho-sites affected by each inhibitor (Fig. 2E). These overlapping targets are highly likely to bona fide targets of the ATR pathway, although not all of them are direct ATR targets. Phospho-motif analysis showed that around half the sites downregulated by both berzosertib or gartisertib conform to the S/T-Q consensus motif typically phosphorylated by ATR and related kinases such as ATM and DNA-PK. However, the majority of remaining sites in common show enrichment for the motif p<u>S</u>RXXS (Figs. 3C, D). LUZP1 Ser570 conforms to the p<u>S</u>RXXS motif, and we showed that its phosphorylation is dependent on ATR but not CHK1 (Fig. 4D-F). We speculate that phosphorylation of these motifs requires an ATR-activated kinase other than CHK1. Candidate kinases include those belonging to the SRPK family of kinases, which target serine residues which lie in SR motifs (88), and this will be interesting to investigate especially as to date these kinases have not been linked to maintenance of genome stability. Intriguingly, enrichment of non-overlapping phospho-motifs specific to each individual inhibitor was observed: YX[pS/pT] for berzosertib and pSXXXTP YXS/T for gartisertib. The most likely explanation is that reduced phosphorylation of these motifs is caused by off-target effects of the inhibitors, but this remains to be investigated.

The first of the three classes of ATR target we defined contain phosphorylation sites that are highly sensitive to ATRi and these are prime candidates for new ATR biomarkers. The phospho-site most sensitive to ATRi is Ser597 of RAP80 (Fig. 2F). ATR was reported to phosphorylate RAP80 at Ser205 (p<u>S</u>Q) after UV exposure (89), but phosphorylation of Ser597, which does not conform to the classical consensus motif for ATR or CHK1, has not been reported previously. In this light, it could be argued that a new ATR biomarker for use in clinical trials should correspond to a classical pS/pT-Q motif. PMS2 pSer603, RAD18 pSer368 and claspin pSer921 all fit this description (Fig. 2F) and therefore represent potentially powerful new biomarkers, especially in trials where ATRi is used as monotherapy.

The second class of new ATR target sites lie in proteins already linked to DDR, and most of these sites lie in pS/pT-Q motifs including FANCI (Ser1120), MCM9 (Ser663) and SMC1 (Ser358) and EXO1 (Ser714) (Table S5). We validated phosphorylation of DHX9 Ser321, which lies close to the helicase domain, and it will be interesting to test if ATR regulates DHX9 activity (Figs. 3A, C). Intriguingly, DHX9 is unusual in that it is excluded from DNA damage sites (Fig. 3F), and this will be interesting to investigate further. A small number of other phosphorylation sites in this class 2 hits lie in functionally annotated domains: for example, within the FANCJ helicase domain (Thr113 and Ser226) and the TLK2 kinase domain (Ser686). Neither protein is known to be regulated by ATR; it will be interesting to test if the catalytic activity of these enzymes is altered in a phospho-dependent manner in response to replication stress or DNA damage. Class 2 targets also include proteins linked to cellular processes known to be controlled by ATR. For example, TERF2IP (RAP1), which binds to the shelterin component TRF2 and plays multiple roles in telomere protein and length maintenance (90). It will be interesting to test the functional impact on telomere biology of mutating the ATR-dependent phosphorylation site in RAP1.

The third class of new ATR target sites lie in proteins not previously linked to DDR or RS responses and they all represent candidate DDR factors. In this light, the majority of the proteins in this class scored positive for recruitment to sites of laser micro-irradiation, consistent with a DDR role. Some of the class 3 proteins have well defined cellular roles. For example, NCAPD2, a subunit of condensin is phosphorylated in an ATR-dependent manner on Ser1310, which is a pSQ site making NCAPD2 likely to be a direct ATR target. This site was identified previously as a pSQ site phosphorylated in response to IR (91), although the kinase responsible was not determined. Here we showed that NCAPD2 is an ATR target and is recruited to DNA damage sites, which points to a DDR role. It will be interesting to investigate the underlying mechanisms and whether the other condensin subunits are recruited. It is tempting to speculate that NCAPD2 recruitment mediates the condensation of chromatin at DNA damage sites that has been linked to genome maintenance (92).

The cellular roles of some of the class three proteins is either unclear or only beginning to be understood. One of these proteins, SUGP1 is also one of the proteins whose phosphorylation is most sensitive to ATRi. SUGP1 is phosphorylated in an ATR- and CHK1-dependent manner on Ser326 and is recruited to DNA damage sites in a manner that depends on PAR synthesis. The role of SUGP1 is only beginning to be understood, but it was recently identified as a binding partner of SF3B1, a spliceosome component that is commonly mutated in cancers and myelodysplastic syndrome. Most of the disease-causing SF3B1 mutations lie in the HEAT repeats, and some cluster in a small area within these repeats including the most frequently mutated residue Lys700 (93). This mutation was shown to disrupt interaction with SUGP1 selectively, and deletion of SUGP1 recapitulates the splicing errors seen in cells expressing the SF3B1 K700 mutation (94). There are already intriguing links between SF3B1 and the DDR – for example, SF3B1-mutated cells accumulate DNA damage and R-loops and show reduced apoptotic responses to DNA damage (95, 96). Moreover SF3B1-mutated MDS cells show increased levels of RS and ATR activation (97). Therefore, it is clear that SUGP1, in complex with SF3B1, is likely to be a key player in DDR. It is possible that SUGP1-SF3B1 impacts on DNA damage responses through controlling the splicing of DDR factors, as was shown recently for DYNLL1 (98). However, the observation that SUGP1 is recruited to DNA damage sites argues that it may have a more direct role, perhaps in collaboration with SF3B1. It will be important to study regulation of this complex by the ATR-dependent phosphorylation. The Ser326 site in SUGP1 conforms to neither the ATR or CHK1 consensus motifs and so we speculate that a CHK1-activated kinase is responsible, but no such kinase is yet known.

*SCAF1* was one of a range of genes identified in a screen for gene deletions that render *BRCA1*-knockout cells resistant to PARP inhibitors (84). Other positives identified in this screen included *53BP1* and components of the shieldin complex; deleting these factors partly restored DNA end resection, RAD51 foci and HR in *BRCA1*-KO cells. Our identification of SCAF1 as an ATR target, reinforced the notion that it may regulate genome stability which we explored further. In this light, SCAF1 depletion or deletion not only reverses the olaparib sensitivity of *BRCA1*-knockout cells, but also causes a modest restoration of HR judged by RAD51 foci (Fig. 6). At present the mechanistic basis for this rescue is unclear. Factors such as shieldin that emerged from the same genetic screen inhibit HR in *BRCA1*-KO cells by suppressing DNA end resection. We found a modest increase in ssDNA levels after IR in *SCAF1*-deleted *BRCA1*-KO cells, judged by BrdU foci, suggesting increased DNA end resection. However, no corresponding increase in levels of phospho-RPA, an alternative readout of resection, was observed. This discrepancy maybe explained by the modest effect size of SCAF1 deletion in restoring RAD51 foci and reversing olaparib sensitivity *BRCA1*-KO cell. The modest increase in resection may not have reached the threshold for triggering RPA phosphorylation. It is interesting to note that deleting *SCAF1* in BRCA1-proficient hTERT *TP53*^−/−^ RPE-1 cells appeared to decrease the levels of ssDNA after IR – the opposite to *BRCA1*-deficient cells. The basis for this surprising observation is not yet clear, but it may be that SCAF1 has different roles in different genetic contexts. Whatever the case, it will be interesting to explore the mechanisms underlying the impact of SCAF1 on HR in *BRCA1*-KO cells, and a key question concerns the relevance of the phospho-dependent interaction of SCAF1 with RNAPII. Given this association, it is possible that SCAF1 impacts HR by influencing the expression - or splicing - of one or more genes involved in control of HR. However, our finding that SCAF1 localizes to DNA damage sites suggests a more direct role. It is possible that SCAF1-regulated transcriptional processes impact on HR, independent of changes in gene expression. Understanding the links between SCAF1 association with RNAPII CTD and its role in inhibiting HR, at least in *BRCA1*-knockout cells will be important.

## Materials and Methods

### Reagents

All the reagents used in the current study including antibodies, siRNA sequences, cDNA clones, oligonucleotides, sgRNA sequences and peptides are listed in Table S7. Full description of the mass spectrometric methods for global phosphoproteomic screening, including sample preparation and data analysis are given in Supplementary Materials.

### Cell lines and cell culture

RPE1 hTERT *TP53^−/−^* Cas9 and RPE1 hTERT *TP53^−/−^ BRCA1 KO* Cas9 cells were a kind gift of D. Durocher (Lunenfeld-Tanenbaum Research Institute, Mount Sinai Hospital, Toronto, Ontario, Canada). All the cell lines were grown at 37°C in a humidified incubator with 5% CO_2_. U-2 OS, U-2 OS Flp-In T-Rex, RPE1 *hTERT* TP53^−/−^ Cas9 and RPE1 *hTERT TP53^−/−^ BRCA1 KO* Cas9 cells were cultured in high glucose Gibco Dulbecco’s modified Eagle’s medium (DMEM, Life Technologies, Paisley, UK) supplemented with 1 mM L-Glutamine, 10% (v/v) fetal bovine serum (FBS), 100 U/mL penicillin, and 100 µg/mL streptomycin. All cell lines were routinely tested and monitored for mycoplasma contamination.

### Cell synchronisation and drug treatment

U-2 OS cells were synchronized in S phase by thymidine-Nocodazole block. Briefly, cells were treated with 2 mM thymidine for 24 h, followed by release into fresh media for 3 h. Cells were then treated with 100 ng/mL nocodazole for a further 12 h. After treatment, cells that remained adherent were gently washed with PBS, released into fresh media for 11 h to enrich for S phase cells. Rounded and floating cells were collected, washed once in PSB and reseeded. S phase cells were either mock treated or treated with 1 µM ATR inhibitors (listed in Table S7) for 1h, followed by replication stress induction with 1 mM hydroxyurea (HU) for 30 minutes. Following HU treatment cells were processed for global phosphoproteomic analysis (detailed in the supplementary Materials & Methods section).

### Cell cycle analysis

Following thymidine-Nocodazole block and release, cells were harvested by trypsinisation at different times post-release. At each time point cells were fixed with ice cold 70% ethanol overnight at −20°C and stained with Propidium Iodide for 30 minutes at room temperature (PI; 50 µg/mL in PBS containing 5% FBS, 0.1 mg/mL RNase A). Cells were analysed by flow cytometry (FACS canto, BD Biosciences) using DIVA software and data analysis to determine cell cycle phases were performed using FlowJo software.

### Cell transfections

For transient expression of GFP-tagged proteins, 1×10^5^ U-2 OS or U-2 OS Flp-In T-Rex cells were seeded in 35 mm glass bottom dishes (FD35-100, WPI); cells were transfected with 1-2 µg of pcDNA5 FRT/TO plasmids containing the gene of interest using GeneJuice transfection reagent (Cat#70967, Merck Millipore) according to manufacturer’s protocol. 8 h post transfection, cells were induced with 1µg /mL tetracycline hydrochloride for 24 h for protein expression. For siRNA mediated protein knockdown, cells were transfected with 50 nM siRNA SMARTpools or individual siRNA using lipofectamine RNAi-Max transfection reagent (13778150, Invitrogen, Paisley, UK) according to manufacturer’s protocol. Cells were analysed or processed for downstream application after 48-72 h of transfection. siRNA sequences and source are provided in Table S7.

### Stable cell lines generation using Flp-In-Trex system

Cells stably expressing GFP-tagged protein of interest were generated as previously described (Khanam *et al*, 2021). Briefly, U-2 OS Flp-In T-Rex cells co-transfected with POG44 Flp-In recombinase expression vector and pcDNA5 FRT/TO - protein of interest in 9:1 ratio, using PEI Max transfection reagent. 48 h post transfection, cells were selected and maintained using the 100 µg/mL hygromycin and 10 µg/mL blasticidin in the medium. After 2 weeks, the surviving colonies were analysed for target protein expression using tetracycline hydrochloride (Cat#T3383; Sigma-Aldrich). Stable cells expressing full length and truncated versions of GFP-SCAF1 were generated in U-2 OS Flp-In T-Rex cells stably expressing mCherry-XRCC1. Briefly, HEK293FT packaging cells were co-transfected with pBabeD-Puro retroviral vector containing mcherry-XRCC1 along with GAG/Pol and VSVG constructs to generate retroviruses, using PEI Max transfection reagent. 48 h post transfection medium containing virion particles were filtered, and target cells were transduced in presence of 8 µg/mL polybrene for 24 h. Cells were selected using fresh media containing 1 µg/mL puromycin. Surviving cells were pooled, single cells with low mCherry-XRCC1 expression were sorted using MA900 multi-application cell sorter (Sony Biotechnology).

### Immunoblotting

For the whole cell extracts, cell pellets were lysed on ice for 30 min in ice-cold RIPA buffer (10 mM Tris-HCl (pH 7.5), 150 mM NaCl, 0.5 mM EDTA, 0.1% SDS, 1% Triton X-100, 1% Sodium deoxycholate, 2.5 mM MgCl_2_) supplemented with protease inhibitor cocktail (cOmplete™ EDTA free protease inhibitor cocktail), phosphatase inhibitor cocktail-2 (Cat#P5726, Sigma-Aldrich) at 1% (v/v), Universal nuclease (Cat#88700, Pierce™ Universal Nuclease) at a final concentration of 250 U/mL, microcystin-LR (Cat#33893, Sigma) at a final concentration of 10 ng/mL with intermittent mixing for every 10 minutes. Lysates were cleared by centrifugation at 17,000 g for 10 min, supernatants were collected for protein estimation by BCA assay. 50 μg of total protein was mixed with quarter of a volume of 4 x LDS sample buffer (Cat#NP0007, Invitrogen™ NuPAGE™ LDS Sample Buffer) and resolved on 4-12% Bis-Tris SDS PAGE gradient gels (NuPAGE, Thermo Fisher). Proteins were electrophoretically transferred onto 0.45 µm nitrocellulose membranes (Cat#10600002, Amersham Protran 0.45 um Nitrocellulose) at 200 mA constant current for 2 h on ice in Tris-Glycine transfer buffer with 20% (v/v) methanol, followed by blocking the membrane with 5% non-fat dry milk in TBS-Tween-20 (0.1% (v/v)) for 30 minutes at room temperature. The blots were probed with respective primary antibody and incubated overnight at 4°C (conditions for each primary antibody used in this study are listed in Table S7). The membrane was washed three times with excess of TBS-Tween-20 (0.1% (v/v)), probed with corresponding secondary antibody diluted in blocking buffer and incubated 1h at room temperature. Blots were washed three times with TBS-Tween-20 (0.1% (v/v)) and once with 1x TBS prior to acquiring bands using LI-COR Odyssey CLx Western Blot imaging system. Primary and secondary antibodies used for Immunoblotting are listed in Table S7.

For detection of ATR activation by immunoblotting using phospho-CHK1/phospho-Rad17 levels in Figure 1 and 2, cells either scraped or cell pellets were lysed directly in 1 x LDS sample buffer (Invitrogen™ NuPAGE™ LDS Sample Buffer, catalogue number: NP0007) supplemented with 2% (v/v) 2-mercaptoethanol and samples were sonicated using a Bioruptor sonicator (Diagenode SA, Seraing, Belgium) at high amplitude for five 30 s on and off cycles. Samples were boiled at 95°C for 5 min before proceeding for immunoblotting as described above.

### SCAF1 gene knockouts in RPE1 hTERT TP53^−/−^ Cas9 and BRCA1 KO Cas9 cells

To knockout SCAF1 from cells, we identified two guide RNA sequences targeting exon 5 of human SCAF1 gene. Single-guide RNA sequences were cloned into px459 GFP plasmid and cells were transfected with 2 µg of plasmid DNA (px459 GFP; sgRNA sequences in Table S7) using PEI Max transfection reagent (Cat#24765-100, Poly sciences). 8 h post transfection, media was replaced with fresh medium and cells allowed to grow for 48 h. Following 48 h of transfection, cells were harvested by trypsinisation, processed for single cell sorting. GFP positive single cells were sorted in 96 well plate using a MA900 multi-application cell sorter (Sony Biotechnology). Single cell clones were maintained in conditioned media with 20% FBS, at 37°C humidified incubator with 5% CO_2_ until visible colonies are formed. Single cell clones of *BRCA1 KO* cells were grown under hypoxic condition with 3% CO_2_. Loss of protein was verified by immunoblotting and immunoprecipitation using SCAF1 polyclonal sheep antibody generated in house at DSTT (DA164, 1^st^ bleed). Individual clones that showed no detectable SCAF1 protein were selected and genomic DNA around the exon five were amplified by PCR (Genotyping Primers Table S7). PCR products were cloned using the StrataClone PCR cloning kit (Cat #240205, Agilent Technologies) and sequenced using T3 and T7 oligonucleotide to confirm absence of wild type allele.

### GFP pulldowns: SCAF1 interaction with RNA Pol II

U-2 OS Flp-In T-Rex cells stably expressing Tet-inducible GFP SCAF1 or GFP-SCAF1 (1187-1312) were induced overnight with 1µg/mL tetracycline hydrochloride. Cells were treated with DMSO or with 10 µM Flavopiridol (Cat#S1230, Selleckchem) or 10 µM THZ1 (Cat#S7549, Selleckchem) for 4 h. Prior to harvest, cells were washed once with PBS and scraped on ice in cold lysis buffer (50 mM Tris-HCl (pH 7.4), 270 mM Sucrose, 150 mM NaCl, 1% (v/v) Triton X-100, 0.5% (v/v) NP-40, protease inhibitor cocktail, phosphatase inhibitor cocktail-2, 10 ng/mL microcystin-LR, benzonase (Novagen, 50 U/mL) and incubated 30 min on ice with intermittent mixing every 10 min by pipetting up and down. After 30 min, lysates were cleared by centrifugation at 17,000 g for 10 min at 4°C. Supernatant was collected, and protein concentration was estimated by the BCA assay. For anti-GFP immunoprecipitations, lysates were pre-cleared with Protein A/G Sepharose beads equilibrated with lysis buffer and incubated for 30 min at 4°C on an end-over wheel. Pre-cleared lysates were used for immunoprecipitations using GFP-trap Sepharose beads (DSTT). Prior to immunoprecipitation, GFP-trap beads were washed twice in lysis buffer and incubated with precleared lysates (2 mg) for 90 min at 4°C on end-over wheel. Beads were washed three times for 3 min with ice cold lysis buffer and a final wash with cold PBS. Immunoprecipitates were denatured by boiling samples in 2x LDS sample buffer supplemented with 2% (v/v) 2-mercaptoethanol at 95°C for 5 min. Proteins were resolved by 4-12% SDS PAGE (NuPAGE) and electrophoretically transferred onto the nitrocellulose membrane. Immunoprecipitates and inputs were analysed by immunoblotting using appropriate primary and corresponding secondary antibodies. Protein bands were acquired using LI-COR Odyssey CLx Western Blot imaging system.

### Peptide pulldown assays

Peptide binding assays were carried out using biotinylated a heptad repeat CTD peptide (Table S7 for peptide sequence) either unphosphorylated or phosphorylated at S2 and S5 of the heptad (pCTD), and bacterial purified MBP or His_6_-tagged SCAF1 SRI domain (1187-end) protein. Briefly, 5 µg of non-phospho or phospho-CTD peptide (per condition) was conjugated to 10 µL (per condition) high-capacity streptavidin agarose beads (Cat#20357, Thermo Scientific) in peptide binding buffer (10 mM Tris-HCl pH 7.5, 150 mM NaCl, 270 mM sucrose, 0.1 mM EGTA, 0.1% (v/v) BME, 0.03% (v/v) Brij 35, protease inhibitor cocktail, phosphatase inhibitor cocktail- 2) for 30 min at room temperature, and washed twice with excess of peptide binding buffer. Beads were then incubated with 3 µg of purified MBP or 6His SCAF1 SRI (1187-end) protein in peptide binding buffer for 90 min at 4°C on a thermomixer with continuous shaking at 1000 rpm. Beads were washed three times with excess of binding buffer followed by a final wash with ice cold PBS. Protein were eluted by boiling beads in 2x LDS sample buffer at 95°C for 5 min and resolved on 4-12% Bis-Tris SDS-PAGE gradient gels Proteins were visualised by staining gel with InstantBlue® Coomassie protein stain (Cat#ab119211; abcam). Lambda phosphatase treatment was carried out as indicated according to the manufacturer’s protocol. Peptide bound beads were washed with 100 µL 1x lambda phosphatase buffer prior to incubating with lambda protein phosphatase (P0753; NEB).

### RAD51 Immunofluorescence (Figs. 6C, E; JR lab)

For RAD51 immunofluorescence in Figs 6C, E, RPE1 hTERT *P53^−/−^* or RPE1 hTERT *P53^−/−^ BRCA1 KO* or *SCAF1 KO* cells were treated with respective siRNA where appropriate using RNAi Max transfection reagent according to manufacturer’s protocol. 24 h post transfection, cells were trypsinised and counted and 5000 cells/well were plated in a Cell star® clear flat bottom 96 well plate (Cat#655090, Greiner bio one) using 6 technical replicates per condition. After 36 h cells were treated with 10 Gy of IR and allowed to recover for different lengths of time (as indicated). At each time point cells were washed once with PBS, fixed with 3% paraformaldehyde in PBS (Santacruz) for 15 min at room temperature followed by two PBS washes and permeabilization with 0.2% triton X-100 in PBS for 5 min at room temperature. Post permeabilization, cells were again washed twice with PBS followed by blocking with blocking buffer (DMEM (Gibco) +10% FBS) for 30 min at room temperature. Cells were co-stained with Rad51 (1:1000) and γH2AX (1:2000) primary antibodies in blocking buffer for 2 h at room temperature followed by three washes with PBS. Cells were incubated with appropriate secondary antibodies diluted in blocking buffer with 1 µg/mL DAPI for 1 h at room temperature. Finally, cells were washed three times with PBS and were left in PBS until image acquisition. Images were acquired and analysed with ScanR High Content Screening Microscopy (Olympus).

### RAD51 immunofluorescence (Fig. 6D; SN lab)

Cells were transfected with siRNAs against SCAF1 using RNAiMax and grown on glass coverslips. 48 hr post-transfection, cells were irradiated with 10 Gy and fixed 3 hr post-IR. Simultaneously, RNA was isolated for qPCR to assess SCAF1 depletion. Cells were fixed for 20 min using 1% PFA, 0.5% Triton X-100 in PBS, followed by another 20 min using 1% (v/v) PFA, 0.3% (v/v) Triton X-100, 0.5% (v/v) methanol in PBS. Cells were blocked in PBS+ (5g/L BSA, 1.5 g/L glycine in PBS) for 30 min at room temperature, followed by primary antibody incubation in PBS+ for 1.5 hr at room temperature (1:15,000 Rb-anti-RAD51 (#70-001, BioAcademia, Japan) and 1:5,000 M-anti-γH2AX (#05-636, Millipore, USA)). Cells were incubated with fluorescently labelled secondary antibodies (1:1,000 G-anti-Rb-AlexaFluor-488, G-anti-M-AlexaFluor-555, ThermoFisher Scientific) and DAPI in PBS+ for 1.5 hr at room temperature before mounting with Aqua-Poly/Mount (Polysciences, USA). Cells were imaged using a Zeiss Axio Imager 2 fluorescent microscope and cells with more than 5 RAD51 foci were manually counted.

### BrdU foci

Cells were incubated with medium containing 10 µM BrdU (Sigma) for 18 h, followed by no treatment or treated with 10 Gy irradiation and 3 h release. Cells were pre-extracted on ice for 8 min using two sequential extraction buffers. Pre-extraction buffer 1 (10 mM PIPES pH 7.0, 300 mM sucrose, 100 mM NaCl, 3 mM MgCl_2_, and 0.5% TritonX-100) and followed by pre-extraction buffer 2 (10 mM Tris-HCl pH 7.5, 10 mM NaCl, 3 mM MgCl_2_, 1% NP-40, and 0.5% sodium deoxycholate). Cells were washed three times with PBS followed by fixation with 4% paraformaldehyde (w/v) for 20 min on ice. After three PBS washes, cells were permeabilized in 0.5% TritonX-100 for 10 min and blocked in 3% BSA-PBS for 20 min on ice. Cells were then incubated with primary antibody against BrdU (1:1000, Fisher Scientific) and PCNA (1:1000, Novus) overnight at 4 °C, followed by three PBS washes and incubation with Alexa Fluor 488 goat anti-mouse (1:1000, Thermo Fisher) and Alexa Fluor 568 goat anti-rabbit (1:1000, Thermo Fisher) secondary antibody for 1 h at room temperature. Coverslips were mounted onto slides with ProLong® Gold Antifade Mountant with DAPI (Invitrogen Life Technologies).

### pRPA (S4 and S8) foci

The protocol was adapted from a previous report (99). Cells were treated with or without 10 Gy irradiation and 3 h release. Cells were then pre-extracted with buffer containing 25 mM HEPES pH 7.9, 300 mM sucrose, 50 mM NaCl, 1 mM EDTA, 3 mM MgCl2, and 0.5% TritonX-100 for 5 min on ice twice. Cells were by fixed by 4% paraformaldehyde for 20 min at room temperature followed by another fixation with methanol for 5 min at −20 °C. After two PBS washes, permeabilization was carried out in 0.5% TritonX-100 for 15 min. After one wash with PBS supplemented with 0.1% tween 20 (0.1% PBST), 2% BSA-PBS was used for blocking for 45 min. Cells were incubated with primary antibody against RPA2-phospho S4+S8 (1:500, Abcam) and PCNA (1:1000, Novus) for 2 h at room temperature, followed by three washes in PBST and incubation with Alexa Fluor 488 goat anti-mouse (1:1000, Thermo Fisher) as well as Alexa Fluor 568 goat anti-rabbit (1:1000, Thermo Fisher) secondary antibody for 1 h at room temperature. Coverslips were mounted onto slides with ProLong® Gold Antifade Mountant with DAPI (Invitrogen life technology).

### Protein extraction and immunoblotting (RPA experiments in Fig. S4F)

Cells were collected and lysed in lysis buffer containing 300 mM NaCl, 1% Triton X-100, 50 mM Tris-HCl pH8, 5 mM EDTA, and 1 mM DTT supplemented with protease inhibitors (1 mM PMSF, 3.4 µg/ml Aprotinin and 1 µg/ml Leupeptin) and phosphatase inhibitors (5 mM NaF and 1 mM Na_3_VO_4_) for 30 min on ice. Samples were sonicated using a Bioruptor sonicator (Diagenode) for 10 cycles (30 s ON/OFF at high power) and centrifugated for 20 min at 4°C. Supernatants were collected and dosed by Bio-Rad Protein Assay Dye Reagent. Equal amounts of total protein were separated by SDS–PAGE and then transferred to nitrocellulose membrane (BioRad) and immunoblotted with antibodies.

### Clonogenic survival assay

sgRNAs targeting SCAF1 (g1: CCACGGACAGCTTCCTCGCA; g3: CTCGGTGTCATGGCCTTCGA) were cloned into pLentiGuide-NLS-GFP as described before (Noordermeer et al., 2018). Viral supernatants were produced using HEK293T cells upon jetPEI-mediated transfection (Polyplus, France) with pLentiGuide-NLS-GFP and third generation packaging vectors. Viral supernatants were collected 48 hr post transfection. RPE1 hTERT *TP53*^−/−^ or *TP53*^−/−^ *BRCA1*^−/−^ cells virally expressing flag-Cas9 (Noordermeer et al., 2018) were transduced with the indicated sgRNAs (or an empty vector as control) and transduced cells were selected using 10 (*TP53*^−/−^) or 15 (*TP53*^−/−^ *BRCA1*^−/−^) μg/mL puromycin for 6 days before seeding cells for clonogenic survival assays. DNA was collected to determine sgRNA targeting efficiency using genomic PCR of the targeting region and TIDE analysis (Brinkman et al., 2014). Targeting efficiencies were 58% for g1 and 70-78% for g3. Alternatively, cells were transfected with siRNAs against SCAF1 (Dharmacon, ON-TARGET plus SMARTpool; Horizon Discoveries, UK) using RNAiMax (ThermoFisher Scientific, USA). Clonogenic survival assays were seeded 48 hr post-transfection. At the time of seeding, RNA was collected for qPCR using a commercially available Taqman primer-probe for SCAF1 (ThermoFisher Scientific, Hs01553675_m1). 250 (*TP53*^−/−^) or 1500 (*TP53*^−/−^ *BRCA1*^−/−^) cells were seeded in 10 cm dishes for clonogenic survival in the presence or absence of 16 nM Olaparib and kept at 37°C, 5% CO_2_ and 3% O_2_. Medium with or without drugs was refreshed 7 days post-seeding. Colonies were stained after 14 days using crystal violet solution (0.4% (w/v) crystal violet, 20% methanol) and manually counted.

### Endogenous SCAF1 immunoprecipitation and mass spectrometry (IP/MS)

SCAF1 WT or KO cells were mock treated or treated with 10 Gy of IR and allowed to recover for 1 h. Endogenous SCAF1 was immunoprecipitated using 10 mg lysate protein using anti-SCAF1 antibody (DA164, DSTT, University of Dundee). Cells were lysed in ice cold lysis buffer (50 mM Tris-HCl (pH 7.4), 270 mM sucrose, 150 mM NaCl, 1% Triton (v/v) X-100, 0.5% (v/v) NP-40, protease inhibitor cocktail, phosphatase inhibitor cocktail-2, 10 ng/mL microcystin-LR, benzonase (Novagen, 50 U/mL) and incubated 30 min on ice with intermittent mixing by pipetting up and down every 10 min. After 30 min, lysates were cleared by centrifugation at 17,000 g for 10 min at 4°C. Supernatant was collected, and protein was estimated by the BCA assay. Samples were pre-cleared by incubating lysates with equilibrated Protein A/G beads for 30 min at 4°C on an end-over wheel. SCAF1 was immunoprecipitated from pre-cleared lysates using sheep polyclonal SCAF1 antibody (1^st^ bleed, DA164, DSTT). Approximately 20 µg of anti-SCAF1 antibody (2 µg/ mg of lysate) was conjugated to 50 µL of Protein A/G beads prior to performing immunoprecipitation. Conjugated antibody-bead complex was incubated with the lysates for overnight at 4°C for pull downs. Immunoprecipitated complex were washed three times with ice cold lysis buffer and finally twice with cold PBS. Samples were boiled at 95°C for 5 min in SDS lysis buffer (5% SDS in 100 mM triethylammonium bicarbonate pH 8.5, complete EDTA free protease inhibitor cocktail (Roche), phosphatase inhibitor cokatail-2, 1 µg/mL microcystin-LR). Eluates from the immunoprecipitates were further processed for S-trap assisted digestion using S-Trap™ micro spin column (Cat#C02-micro-40, Protifi) according to manufacturer’s protocol followed by TMT labelling as described previously for the global phosphoproteomics screen (without phospho-peptide enrichment)(100). After TMT labelling, the quenched samples were mixed and fractionated with high pH reverse-phase C18 chromatography using the Ultimate 3000 high-pressure liquid chromatography system (Dionex) at a flow rate of 500 µl/min using two buffers: buffer A (5 mM ammonium formate, pH 10) and buffer B (80% ACN, 5 mM ammonium formate, pH 10). Briefly, the TMT-labelled samples were resuspended in 200 µl of buffer A (5 mM ammonium formate, pH10) and desalted then fractionated on a C18 reverse-phase column (4.6 × 250 mm, 3.5 µm, Waters) with a gradient as follows: 3% Buffer B for 19 min at 275 uL/min (desalting phase), ramping from 275 uL/min to 500 uL/min in 1 min, 3% to 12% buffer B in 1 min, 12% to 40% buffer B in 30 min, 40% B to 60% B in 5 min, 60% B to 95% B in 2 min, 95% for 3 min, ramping to 3% B in 1 min and then 3% for 9 min. A total of 96 fractions were collected and then concatenated into 24 fractions, which were further speed vacuum-dried prior to LC–MS/MS analysis. Peptides were resuspended in 5% formic acid in water and injected on an UltiMate 3000 RSLCnano System coupled to an Orbitrap Fusion Lumos Tribrid Mass Spectrometer (Thermo Scientific). Peptides were loaded on an Acclaim PepMap trap column (Thermo Scientific #164750) prior analysis on a PepMap RSLC C18 analytical column (Thermo Scientific #ES903) and eluted on a 120 min linear gradient from 3 to 35% Buffer B (Buffer A: 0.1% formic acid in water, Buffer B: 0.08% formic acid in 80:20 acetonitrile:water (v:v)). Eluted peptides were then analysed by the mass spectrometer operating in Synchronous Precursor Selection mode using a cycle time of 3s. MS1 were acquired at a resolution of 120000 with an AGC target of 100% and a maximum injection time of 50 ms. Peptides were then selected for MS2 fragmentation using CID with an isolation width of 0.7 Th, NCE of 35%, AGC of 100% and maximum injection time of 50 ms using the “rapid” scan rate. Up to 10 fragments were then selected for MS3 fragmentation using HCD with an isolation width of 3 Th, NCE of 65%, AGC of 200% and maximum injection time of 105 ms and spectra were acquired at a resolution of 50000. Dynamic exclusion was set to 60 s with a tolerance of +/- 10 ppm. Mass spectrometry raw data was searched using MaxQuant (version 2.1.3.0) (101) against a *homo sapiens* FASTA (42,390 entries, downloaded 18^th^ August 2022, inclusive protein isoforms) from Uniprot (www.uniprot.org). Additionally, to the standard MaxQuant search parameters, deamidation of N and Q was set as variable modification. Data was analysed using R (version 4.1.1) (102) with in-house developed scripts based on previous versions published (100). In brief, intensities of peptides repeatedly measured within a single fraction were averaged. Data was then transformed and calibrated using VSN (103, 104) The median peptide intensities belonging to each respective protein was taken as heuristic for total protein intensity and data were statistically tested using limma (105)(106). Proteins underwent volcano plot analysis and proteins clustering within a group distinct from the bulk of data (107) were regarded as statistically significant (adjusted p-value < 0.08). Mass spectrometry raw data, MaxQuant search parameters and output file, and FASTA file have been deposited at jPOSTrepo (108) and can be downloaded via ProteomeXchange (109) (PXD041201). All data analysis scripts can be downloaded from Zenodo (110) via https://doi.org/10.5281/zenodo.7784982.

### Extracted ion chromatography (XIC) analysis

U-2 OS cells were seeded at 25% confluency in 100 mm plates. 24 h post plating, cells were transiently transfected with plasmid DNA containing GFP tagged protein of interest using PEI Max transfection reagent (24765; Polysciences). Briefly, 3 μg of plasmid DNA and 9 μg of PEI (1:3 ratio of DNA:PEI) were diluted in 1 mL of Opti-MEM reduced serum media, pulse vortexed for 15 sec and the mixture was incubated at room temperature for 15 min. The transfection mixture was then added drop wise to the target cells, 8 h post transfection fresh media was replaced, and cells were incubated further for 24 h. After 24 h, cells were either mock treated or treated with 1 µM gartisertib or 0.5 µM CHK1 inhibitor PF477736 for 1 h followed by hydroxyurea treatment at a final concentration of 1 mM for 30 min. Cells were then harvested by trypsinisation and washed once with ice cold PBS. Cell pellets were lysed in 300 µL of RIPA buffer (10 mM Tris-HCl (pH 7.5), 150 mM NaCl, 0.5 mM EDTA, 0.1% (v/v) SDS, 1% (v/v) triton X-100, 1% (v/v) sodium deoxycholate, 2.5 mM MgCl_2_, Protease inhibitor cocktail, phosphatase inhibitor cocktail-2, 10 ng/mL microcystin-LR) on ice for 30 min and then diluted with 450 µL of dilution buffer (10 mM Tris-HCl (pH 7.5), 150 mM NaCl, 0.5 mM EDTA, Protease inhibitor cocktail, phosphatase inhibitor cocktail-2), incubated at 4°C on end-over wheel for further 10 min. Lysates were then clarified by centrifugation at 13,300 rpm for 10 min at 4°C. An aliquot of supernatant was saved for immunoblot analysis.

For extracted ion chromatography (XIC) analysis, 25 µL of equilibrated GFP-trap Sepharose beads (DSTT, University of Dundee) were incubated with lysates for 90 min at 4°C. Precipitates were washed three times with 1 mL ice cold washing buffer (4 parts of RIPA buffer mixed with 6 parts of dilution buffer). Samples were denatured in 2x LDS sample buffer supplemented with 2% (v/v) 2-mercaptoethanol at 95°C for 5 min. The experiment was carried out in triplicates using lysates from independent replicates per condition. Denatured samples were resolved in 4-12% Bis-Tris SDS-PAGE gradient gels and stained with InstantBlue® Coomassie protein stain (Cat#ab119211, abcam). Protein bands were excised from gel, cut into approximately 1 mm^2^ pieces, and destined using 25 mM AmBiC, 30% (v/v) acetonitrile solution. Proteins were digested with trypsin/LysC protease (Cat#A40009; Thermo Scientific) and labelled with TMT 10plex as described previously (100). Data analysis protocol is described in detail in Supplementary Materials and Methods. The mass spectrometry raw data for LUZP1 and SUGP1 was uploaded to ProteomeXchange via jPOSTrepo can be downloaded from with the identifiers PXD040476, while DHX9 data was uploaded via PRIDE (111) and can be accessed under PXD041250. The data analysis scripts are available via Zenodo (https://doi.org/10.5281/zenodo.7661023 (LUZP1, SUGP1) and https://doi.org/10.5281/zenodo.7788447 (DHX9)).

### Laser micro-irradiation

Recruitment of protein to the site of DNA damage were monitored by laser micro-irradiation as described previously (107) with the following changes. Around 1 x 10^5^ U-2 OS Flp-In T-Rex cells were seeded in 3.5 cm glass bottom dishes (FD35-100) and transiently transfected with the plasmid DNA containing GFP-tagged protein of interest (See Table S7 for plasmid constructs) using GeneJuice transfection reagent (70967, Merck) according to manufacturer’s protocol. After 24 h, the media was changed to complete DMEM containing 10 µM bromodeoxyuridine (BrdU) for another 24 h. Shortly before irradiation, media was replenished with warm phenol-red free media (31053; Thermo Fisher). Cells were placed in a 37°C chamber incubator supplemented with 5% CO_2_ mounted on Leica TCS SP8X microscope system (Leica Microsystems).

For SCAF1 recruitment, cells stably expressing GFP-SCAF1 were seeded in 3.5 cm glass bottom dish in presence of 10 µM BrdU and 1 µg/mL tetracycline hydrochloride for 24 h. Prior to irradiation cells were either mock treated or treated with flavopiridol, THZ1, DRB, olaparib or PDD00017273 (PARGi) as indicated, and cells were placed on chamber incubator at 37°C with 5% CO_2_ attached to an Axio Observer Z1 spinning disc confocal microscope (Zeiss). Image acquisition and image analysis were performed as describe in Khanam *et al* (2021).

## Acknowledgements

We thank the technical support of the MRC-PPU including the DNA Sequencing Service, Tissue Culture team, Reagents and Services team, and the PPU Mass Spectrometry team. We’re grateful to Dan Durocher for kindly providing RPE-1 *hTERT TP53*^−/−^ and RPE-1 *hTERT BRCA1*^−/−^ *TP53*^−/−^ cells, and for useful discussions. Luis Sanchez-Pulido and Chris Ponting for help with bioinformatic analyses. We thank Paul Appleton for help with microscopy. We are grateful to Ulrich Pehl and Claudio Lademann from the healthcare business of Merck KGaA, Darmstadt (Germany) and members of the Rouse team for useful discussions and help with designing the screening experiments. This work was supported by the Medical Research Council (grant number MC_UU_12016/1; MG, PL, FL, IM, JR) and by the healthcare business of Merck KGaA, Darmstadt, Germany (CrossRef Funder ID: 10.13039/100009945) who provided berzosertib and gartisertib free of charge. The work was also supported by Fondation du CHU de Québec and FRQS Ph.D. scholarships (YG and JYM). J.Y.M. is a Canada Research Chair in DNA repair and Cancer Therapeutics. Research in the Alabert lab is funded by the European Research Council ERC-Stg-IDRE.

## Supplementary Materials and Methods

### Phosphoproteomic screening

#### Sample preparation for phosphoproteomic analysis

Five 150 mm plates were seeded with U2 O-S cells at 30% confluency and synchronised to enrich cells in S phase as described in the method section. Cells were mock treated or treated with 1 µM ATR inhibitors (berzosertib or gartisertib) followed by treatment with 1 mM hydroxyurea for 30 min. Post drug treatment, cells were harvested by trypsinisation and washed once with ice cold PBS. To the cell pellets 1.5 mL ice cold TCA-Acetone solution (20 % (v/v) TCA, 80% (v/v) acetone, 0.2% DTT (wt/v)) was added, cells were vortexed vigorously to disperse the pellet and kept overnight at −20°C. After 24 h, samples were centrifuged at 20,000 g at −9°C for 20 min, and the supernatants was discarded. Pellets were washed once with 2 mL ice cold 80% (v/v) acetone, vortexed vigorously, centrifuged again at 20,000 g at −9°C for 20 min. Carefully removed the supernatant and pellets were air dried for 10 min at room temperature. Cell pellets from each plate were then lysed in 350 µL of 8 M urea, 50 mM AmBiC, 1% (v/v) phosphatase inhibitor cocktail-2, 0.1% (v/v) microcystin-LR, pH 8. Samples were mixed thoroughly by pipetting up and down and incubated at room temperature for 15 min. Post incubation samples were sonicated using a Bioruptor sonicator at high amplitude for 10 cycles (30 sec on/30 sec off). Lysates were clarified by centrifugation at 17,000 g for 10 min and samples were stored at −80°C until further use for mass spectrometric analysis. Five independent biological replicates per condition were prepared on different days using two different ATR inhibitors-berzosertib and gartisertib. Protein concentrations were determined using BCA assay.

A total of 3.5 mg total protein from each sample was taken and volume was equalized using 8M Urea (Cat. No. 1.08487.0500, Merck, Darmstadt, Germany), 50mM ammonium bicarbonate (AmBiC, Cat. No. 09830, MilliporeSigma, St. Louis, Missouri, US), 1% (v/v) Phosphatase Inhibitor Cocktail 2 (Cat. No. P5726, MilliporeSigma) and 0.1% (v/v) Microcystin-LR (Cat. No. 33893, MilliporeSigma). Proteins were then reduced and alkylated with 10mM tris(2-carboxyethyl)phosphine (TCEP, Cat. No. C4706, MilliporeSigma), 25mM Chloroacetamide (Cat. No. C0267, MilliporeSigma) in the dark for 45 minutes at room-temperature (RT). Afterwards, the samples were diluted with 50mM AmBiC (MilliporeSigma) to a concentration of 2M Urea. Trypsin/LysC (Cat. No. V5071, Promega, Madison, Wisconsin, US) was added at a 1:50 enzyme:protein ratio and the samples were incubated at 37°C overnight. Samples were acidified by adding trifluoroacetic acid (TFA, Cat. No. 1.08262.0025, Merck) to a concentration of 1% (v/v) and centrifuged at 3,270 x g for 15 minutes at RT. Supernatant was desalted by C_18_ chromatography using Sep-Pak cartridges (830 mg, Cat. No. WAT023635, Waters, Milford, Massachusetts, US), eluted peptides were snap frozen and freeze dried overnight.

#### Phosphopeptide enrichment and TMT labelling

Samples were reconstituted in 300 µL 250mM lactic acid (Cat. No. 252476, MilliporeSigma), 70% (v/v) Acetonitrile (ACN, Cat. No. 1.00029.1000, Merck), 3% (v/v) TFA (Merck). Peptides were incubated with TiO_2_ beads (Cat. No. 5010-21315, GL Sciences, Tokyo, Japan) at a ratio of 1mg peptide : 4mg beads. Beads were washed with 250mM lactic acid, 70% (v/v) ACN, 3% (v/v) TFA, then with 70% (v/v) ACN, 3% TFA and as last step with 0.1% (v/v) TFA. Phosphopeptides were eluted from the beads using 1% (v/v) NH_4_OH (in case of samples derived from berzosertib experimental series: 50% (v/v) ACN (Merck) was added to elution buffer as well), snap frozen and freeze dried overnight. This phosphopeptide enrichment procedure was repeated for a total of 2 times. Phosphopeptide enrichment was checked via LC-MS/MS and an average of 8.66% (gartisertib) and 10.93% (berzosertib) of non-phosphorylated peptides were detected. Subsequently, the phosphopeptides were labelled via tandem mass tags (TMT10plex, Cat. No. 90113, Thermo Fisher Scientific, Waltham, Massachusetts, US). Freeze dried peptides were reconstituted in 100mM Triethylammonium bicarbonate (TEAB, Cat. No. 90114, Thermo Fisher Scientific). TMT labels were resuspended in anhydrous ACN (Cat. No. 271004, MilliporeSigma), added to the respective samples and incubated 2h at RT. In case of the (HU vs. HU + gartisertib) experimental series, the HU group was labelled with TMT126-TMT128C, the inhibitor treatment group was labelled with TMT129N-TMT131. In the experimental series (HU vs HU + berzosertib), the inhibitor treatment group was labelled with TMT126-TMT128C, while the HU group was labelled with TMT129N-TMT131. An aliquot was taken (the remainder samples were interim stored at −80°C) and labelling efficiency was checked via LC-MS/MS. Peptide N-termini were detected as labelled in 96.8% (gartisertib) or 87.2% (berzosertib) of all peptides, and lysines were detected as labelled in 97.8% (gartisertib) or 85.1% (berzosertib) of all cases. The labelling reaction of the remainder samples was stopped using hydroxylamine (Cat. No. 467804, MilliporeSigma). Afterwards, the samples were pooled, snap frozen and freeze dried.

#### Sample prefractionation using high pH reverse phase liquid chromatography

The pooled sample was fractionated using high pH reverse phase liquid chromatography. Peptides were reconstituted in 200 µL 10mM ammonium formate (Cat. No. 70221, MilliporeSigma) pH 10 and separated on a C_18_ column (4.6 x 250 mm, 3.5µm particle size, Cat. No. 186003570, Waters) using a two-component buffer system. Buffer A consisted of 10mM ammonium formate, pH 10 while Buffer B consisted of 80% acetonitrile and 10mM ammonium formate, pH10. Peptide separation was achieved using a 75min gradient with following settings: 0-5.5 minutes 3% (v/v) Buffer B (flowrate: 275µl/min); 5.5-10 minutes linear gradient to 10% (v/v) Buffer B (flowrate: 569 µl/min); 10-45 minutes linear gradient to 35% (v/v) Buffer B, 45-55 minutes linear gradient to 50% (v/v) Buffer B; 55-65 minutes linear gradient to 80% (v/v) Buffer B; 65-67.5 minutes linear gradient to 100% Buffer B; 67.5-70 minutes 100% Buffer B; 70-70.5 minutes gradient to 3% (v/v) Buffer B; 70.5-75 minutes 3% (v/v) Buffer B. 75 fractions were collected, concatenated into 20 fractions, snap frozen, freeze dried and stored at −80°C until further usage.

#### LC-MS/MS

Before LC-MS/MS analysis, samples were reconstituted in 2% (v/v) ACN (Merck), 0.1% formic acid (FA, Cat. No. 5.33002.0050, Merck). LC-MS/MS was performed using a Thermo Dionex Ultimate 3000RLSC system coupled to an Orbitrap Fusion Tribrid mass spectrometer (Thermo Fisher). Samples were injected onto a C_18_ trap column (Acclaim^TM^ PepMap^TM^ 100, Cat. No. DX164564, Thermo Fisher) and washed for 5 minutes using 3% (v/v) ACN, 0.5% (v/v) TFA at a flowrate of 5 µL/min. The column oven was set to 45°C. Peptides were separated on a C_18_ column (EASY-Spray^TM^, 50 cm length, Cat. No. 03-251-874, Thermo Fisher) using a segmented linear gradient applying a two-buffer system with a total run-time of 180 minutes at a flowrate of 300 nL/min. Buffer A consisted of 3% (v/v) ACN, 0.1% (v/v) FA, Buffer B of 80% ACN, 0.08% (v/v) FA. During the first 145 minutes Buffer B percentage increased from 5 to 25%, followed by an increase to 35% B until minute 155, then increase to 95% B until minute 160 with isocratic elution until minute 165, this was followed by a decrease in Buffer B to 3% over 30 seconds with isocratic column wash until minute 180. Eluted peptides were injected into the Orbitrap MS. The ion source spray voltage was operated in positive mode and set to 2000 V. Ion transfer tube was heated to 275 °C. The mass spectrometer was operated in data-dependent Top20 mode. MS full scan was performed using the Orbitrap detector, detecting positively charged ions between 375-1500 *m/z* at a resolution of 120K at 200 *m/z*. AGC target was set to 4 x 10^5^ with a maximal ion injection time of 50 ms. For MS2 scan, precursor ions had to exhibit the following properties: Intensity above 2 x 10^4^, charge states between 2 and 7, while exhibiting an isotopic distribution expected for a peptide. Precursor ions were isolated using a width of 1.6 *m/z* and fragmented by higher-energy collisional dissociation (HCD) with 37% normalized collision energy. AGC target was set to 5 x 10^4^. In case of samples from gartisertib experimental series the maximum injection time was set to 120 ms, while for samples from the berzosertib series this was set to 200 ms. MS/MS spectra were recorded in the Orbitrap detector at a resolution of 50K (at 200 *m/z*). To avoid repeated fragmentation of the same precursor ion species, dynamic exclusion was set to 40 seconds with a mass tolerance of 10 ppm.

#### Data analysis global phosphoproteomics

Mass spectrometry raw data was searched against a *homo sapiens* FASTA file (42,326 entries including protein isoforms, downloaded from www.uniprot.org on 5^th^ April 2018) (1) using MaxQuant (Version 1.6.2.3)(2). Standard search settings were applied with following additional variable peptide modifications: Phosphorylation of serine, threonine and tyrosine, deamidation of asparagine. Peptide false discovery rate (FDR) was set to 5%. All mass spectrometry raw data, the MaxQuant search parameters and output files were uploaded to jPOSTrepo (3) and can be downloaded via ProteomeXchange (4) with the identifier PXD040469. Results were analysed using an in-house developed R (version 4.1.1 (5) computational pipeline, based on modified scripts applied in (6). In brief, within each fraction the peptide intensities were averaged. Data were normalized using variance stabilizing normalization (VSN) (7, 8) and statistically tested using linear models for microarrays (*limma*) (9, 10). A peptide was regarded as differentially abundant between the two conditions if the adjusted p-value was ≤ 0.05. Singly phosphorylated peptides, lower abundant under inhibitor treatment, exhibiting a phosphorylation PTM score probability ≥ 0.994 (corresponding to a false localization rate of 1% (11)) underwent phosphorylation sequence motif analysis using rmotifx (12). GO term analysis was based on a modified version of a previous R script file (6), applying a Fisher’s exact test. For this, the unique leading razor protein names connected to phosphopeptides lower abundant under inhibitor treatment were used as foreground. The unique leading razor protein names of the whole dataset were used as background. Resulting p-values were converted into q-values (13) and GO terms with q ≤ 0.01 and at least 10 protein hits were regarded as enriched in the foreground. Peptide and protein metadata were mined from following databases: Protein interaction partners: BIOGRID (version 4.4.217)(14), phosphorylation sites: PhosphoSitePlus® (version 122022, downloaded from www.phosphosite.org the 20^th^ December 2022)(15), Gene ontology terms: downloaded from www.uniprot.org on 6^th^ January 2023, Protein modifications dataset was downloaded from www.uniprot.org on 6^th^ January 2023. Protein domains data were extracted from Ensembl (16), Interpro (17), Prosite (18) and Pfam (19) using the R biomaRt library (20, 21). List of kinases (pkinfam) was downloaded on 06/05/2020 from https://www.uniprot.org/docs/pkinfam, list of E3 ligases from https://hpcwebapps.cit.nih.gov/ESBL/Database/E3-ligases/ on 28/08/2019 (22), and the list of deubiquitylases (dubs.txt) was downloaded 06/05/2020 from https://www.genenames.org/data/genegroup/#!/group/996. Additional R libraries used were: miscTools (23), ggplot2 (24), reshape2 (25), seqinr (26), plyr (27), GO.db (28), stringr (29), ggrepel (30), ggpointdensity (31), extrafont (32), scales (33), ggseqlogo (34), viridislite (35), xml2 (36), magrittr (37), dplyr (38) and readxl (39). All data analysis scripts are available at Zenodo (40) under https://doi.org/10.5281/zenodo.7784890. The R library version numbers and the R session information is stored in the file “Text_File_S1_Session_Info_Phospho-TMT.txt” that can be accessed under the same URL.

#### Extracted ion chromatogram (XIC) analysis

Protein extraction from SDS-PAGE bands, digestion with Trypsin/LysC followed the protocol described in (41), deviating in the usage of 5% hydroxylamine to stop the TMT labelling reaction. The samples were analysed on an Orbitrap Lumos (Thermo Fisher), connected to a Thermo Dionex Ultimate 3000RLSC system. The identical MS instrument settings and LC gradient as detailed in (6) were applied. Mass spectrometry raw data was searched using MaxQuant (Version 2.1.3.0) using the same FASTA file as described under “*Data analysis global phosphoproteomics*”. Standard MaxQuant search settings were applied with the additional variable modification of phosphorylation of serine, threonine and tyrosine. Peptide FDR threshold was set to 5%. The TMT intensity data for the LUZP1 phosphopeptide AIGALAS<u>pS</u>R (Spectrum #17044 in file LUM190801-16_FW052_TL-01_FW.raw) had to be extracted manually using Xcalibur QualBrowser (Version 2.2 SP1.48, Thermo Fisher) as MaxQuant did not detect these signals although they are clearly present in the raw data. Data were analysed using modified versions of the R scripts used in (6, 41). In summary, data were normalized using VSN and the TMT reporter intensities were statistically tested using t-tests under application of Bonferroni correction of the significance threshold to α = 0.05 / 3 = 0.0167 (3 t-tests). All mass spectrometry raw data, MaxQuant search settings and output files were uploaded to jPOSTrepo and can be downloaded via ProteomeXchange with the identifier PXD040476. Data analysis scripts and session information file can be downloaded from Zenodo under https://doi.org/10.5281/zenodo.7661023. Samples from DHX9 immunoprecipitation were analysed after SDS-PAGE, digestion and peptide extraction as follows: Peptides were resuspended in 5% formic acid in water and injected on an UltiMate 3000 RSLCnano System coupled to an Orbitrap Exploris 240 (Thermo Scientific). Peptides were loaded on an Acclaim PepMap trap column (Thermo Scientific #164750) prior analysis on a PepMap RSLC C18 analytical column (Thermo Scientific #ES903) and eluted on a 60 min linear gradient from 3 to 35% Buffer B (Buffer A: 0.1% formic acid in water, Buffer B: 0.08% formic acid in 80:20 acetonitrile:water (v:v)). Eluted peptides were then analysed by the mass spectrometer operating in data dependent acquisition mode using a cycle time of 2 s. MS1 were acquired at a resolution of 60000 with an AGC target of 300% and a maximum injection time of 25 ms. Peptides were then selected for HCD fragmentation using an isolation width of 1.2 Th, NCE of 30%, AGC of 100% and maximum injection time of 100 ms and MS2 were acquired at a resolution of 15000. Dynamic exclusion was set to 30 s with a tolerance of +/- 10 ppm. Mass spectrometry raw data was searched using MaxQuant (Version 2.2.0.0) against a Uniprot *homo sapiens* FASTA (42,397 entries inclusive isoforms, downloaded 2^nd^ January 2023 from www.uniprot.org). In addition to the standard MaxQuant search parameter, deamidation of N and Q, and phosphorylation of S, T and Y was added as variable modifications. Intensities of peptides from DHX9 which were repeatedly measured were averaged using their median intensity and then all DHX9 peptides from the different samples were normalized via VSN. Data was then statistically tested using limma. The mass spectrometry raw data, MaxQuant search settings and output files were uploaded to PRIDE (42) and can be accessed via ProteomeXchange (PXD041250). Data analysis scripts were uploaded to Zenodo and can be downloaded via https://doi.org/10.5281/zenodo.7788447.

## Supplementary Figure Legends

**Figure S1.**
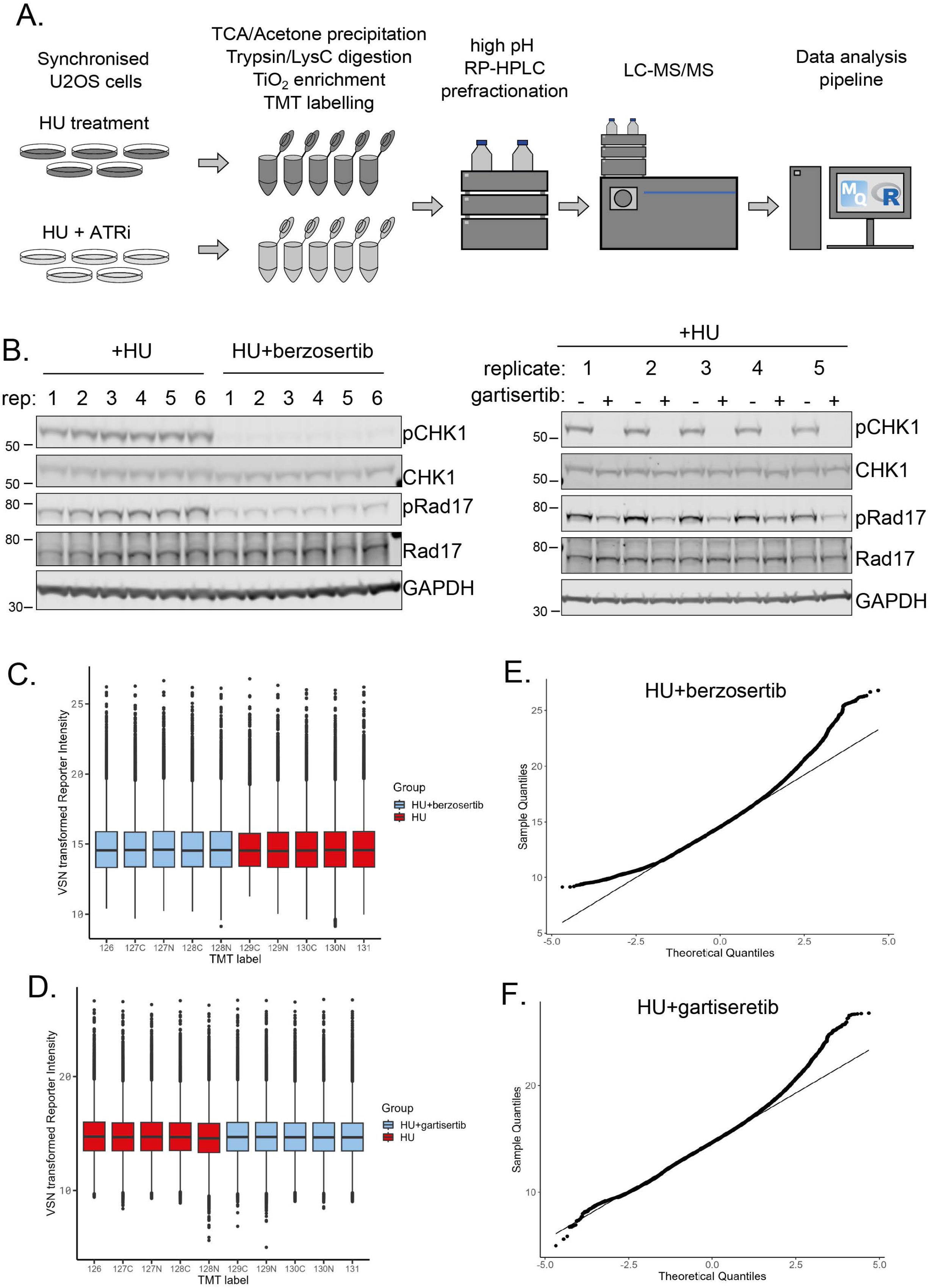
Quantitative phosphoproteomic screening workflow. A. Quantitative phosphoproteomic screening workflow. B. Assessment of ATR activity in the 5 biological replicates used in the HU±berzosertib and HU±gartisertib phosphoproteomic screens. U2-O-S cells synchronized in S-phase (11h after release from nocodazole) were pre-incubated for 1h, or not, with berzosertib or gartisertib (1 μM) before addition of HU (0.5 mM) for 0.5h. Cell extracts were subjected to western blotting with the antibodies indicated. C, D. Boxplot of intensity distribution in each TMT channel (C, HU±berzosertib screen; D, HU±gartisertib screen). No obvious discrepancy between the median values of the individual channels indicates a successful calibration by VSN and no introduction of an obvious intensity bias for any experimental group. E, F. Normal QQ-Plots of the TMT intensity data from the HU±berzosertib screen (E) and HU±gartisertib screen (F) after VSN transformation. Only minor deviations from the line indicates the transformed data follows a normal distribution to a satisfactory degree. The hypervariable datapoints in the upper quantiles are controlled by the application of the robust implementation of the empirical Bayes algorithm used by limma (9) and implemented in the analysis scripts.

**Figure S2.**
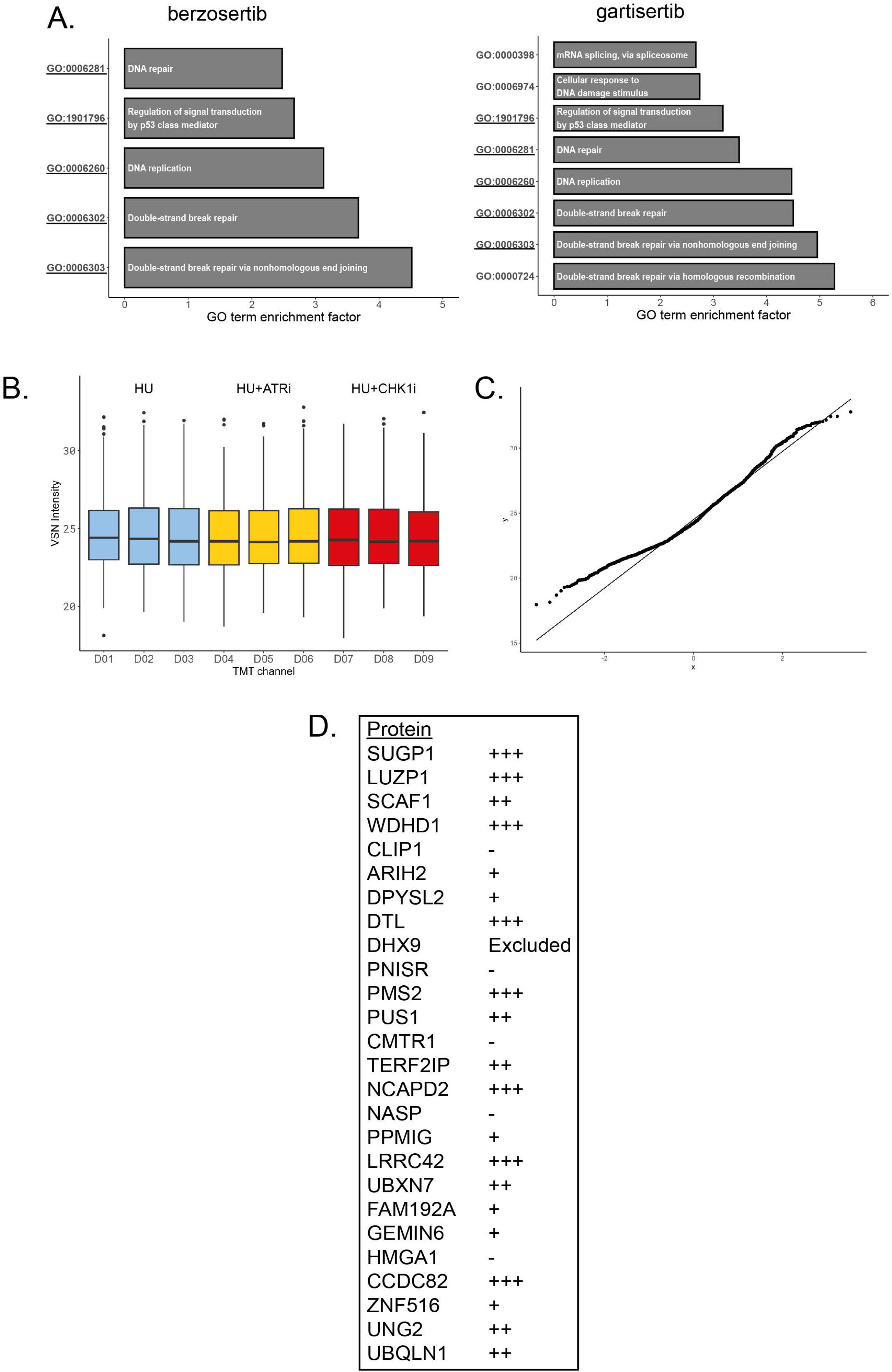
Analysis of Gene Ontology and protein recruitment. A. Gene Ontology terms enriched among proteins whose phosphorylation was inhibited by berzosertib or gartisertib. Significance cut–off was set as α = 0.01 with at least 2 proteins identified in the respective group. B. Relative strength of protein recruitment to DNA damage sites in laser micro-irradiation experiments.

**Figure S3.**
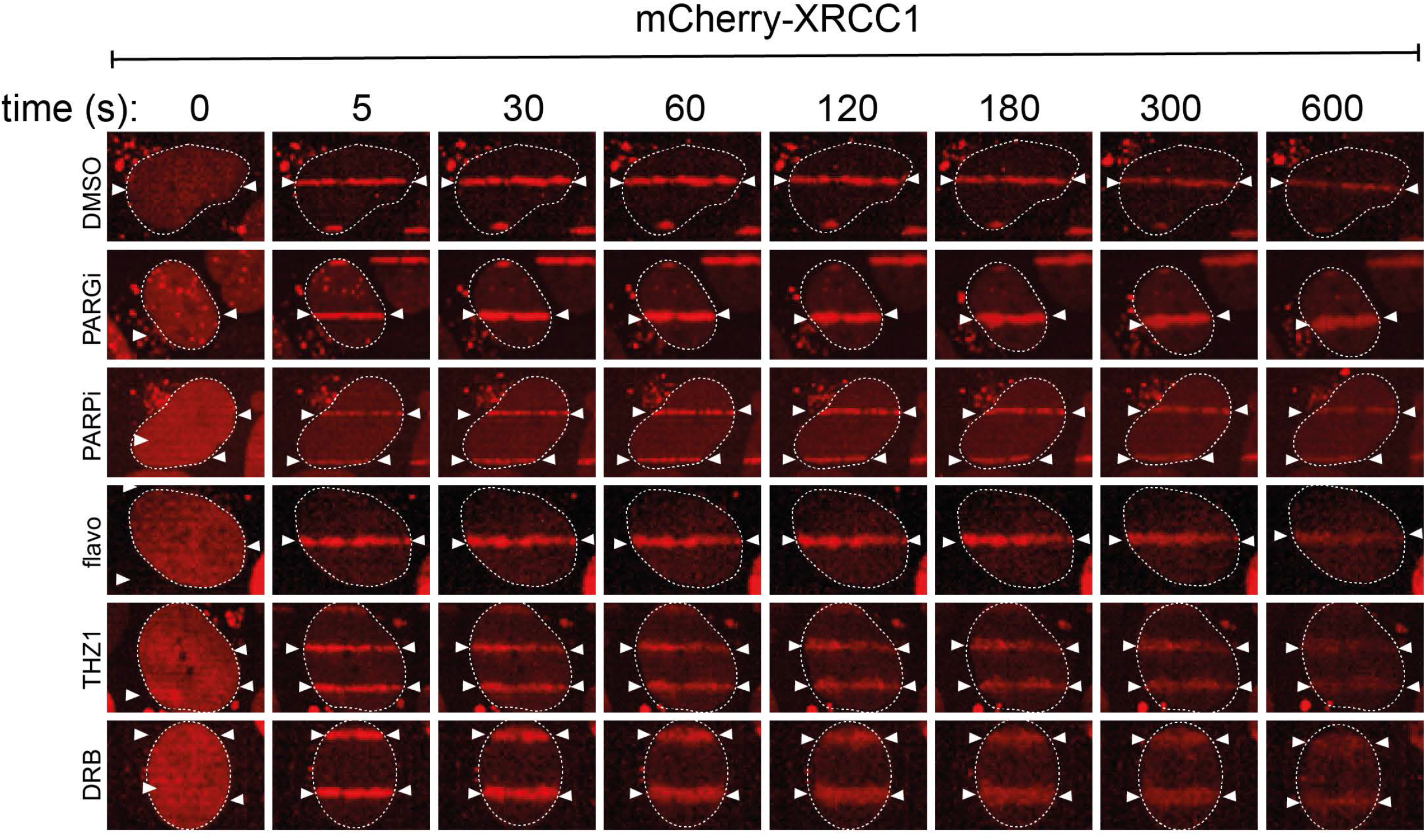
Transcriptional inhibitors do not affect XRCC1 recruitment to DNA damage sites. BrdU–sensitized U–2–OS Flp–In T–REx cells stably expressing mCherry-tagged XRCC1, pre–incubated with DMSO, olaparib (5 μM; PARPi), PDD00017273 (0.3 μM; PARGi), flavopiridol (10μM), THZ-1 (10 μM) or DRB for 1 h were micro-irradiated with a 405 nm laser and imaged at the times indicated.

**Figure S4.**
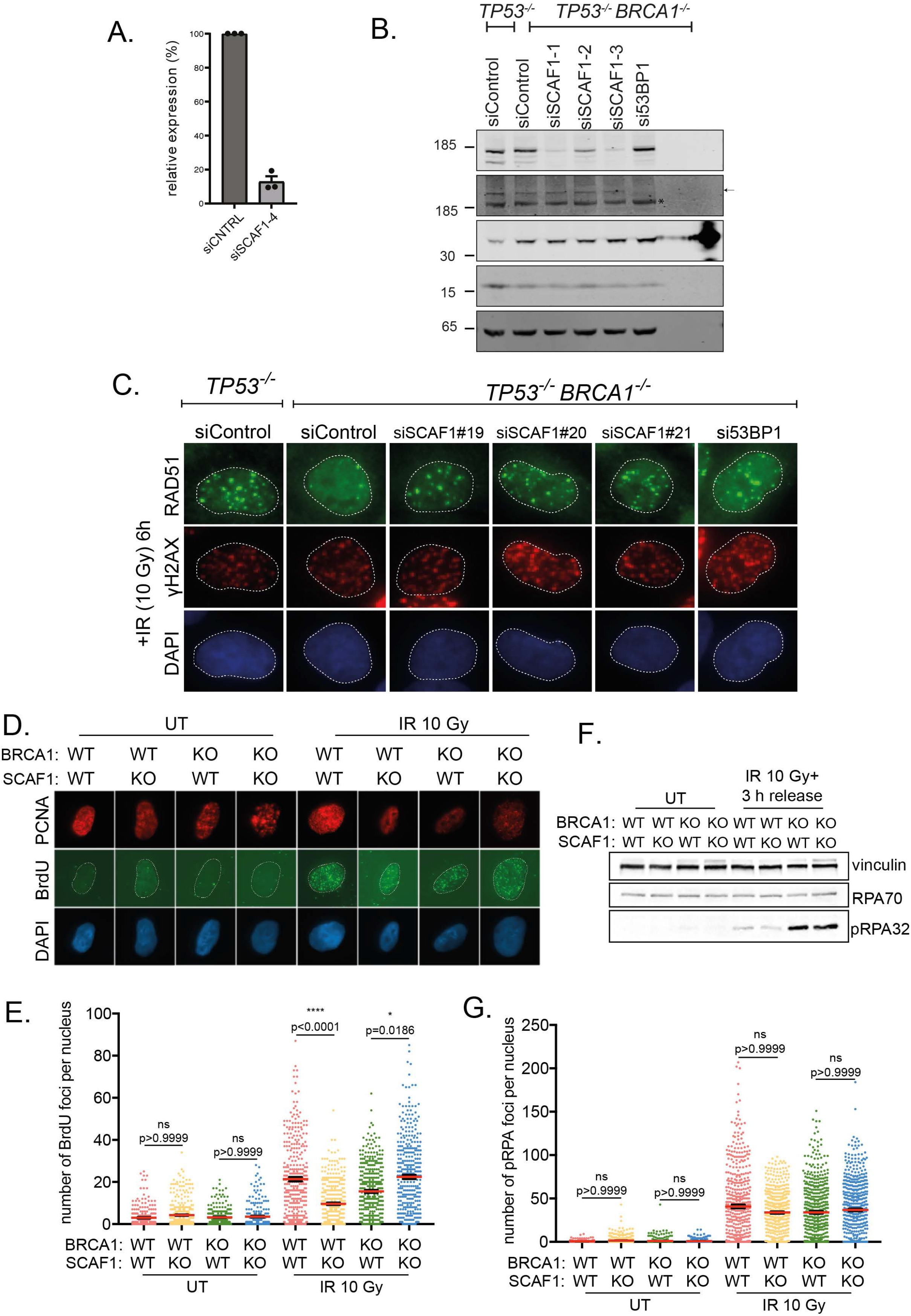
SCAF1 depletion restores RAD1 foci in BRCA1-knockout cells. A. qPCR analysis of SCAF1 knock-down upon siRNA transfection of RPE1 hTERT *TP53^−/−^ BRCA1^−/−^* cells with siSCAF1-4 (SN lab). SCAF1 expression was normalized to expression of the house keeping gene GUSB and normalized expression of SCAF1 in siCNTRL samples was set at 100% per biological replicate (n=3). B. The cell lines shown were transfected with the siRNAs indicated. Cells were lysed and extracts subjected to western blotting with the antibodies indicated. C. The cell lines indicated were transfected with the siRNAs indicated and after 24h cells were exposed to IR (10 Gy). Cells were allowed to recover for 3h, fixed and subjected to immunofluorescence to detect RAD51 foci. D. Representative images of immunofluorescence staining against BrdU and PCNA in RPE1 hTERT *TP53^−/−^* that are wild type (WT) or knockout (KO) for BRCA1 and/or SCAF1 as indicated. E. Quantification of the number of BrdU foci in S-phase RPE1 hTERT *TP53^−/−^* cells with the *BRCA1* and *SCAF1* genotypes indicated: WT, wildtype, KO, knockout determined by PCNA staining before and after exposure of cells to ionising radiation (IR; 10 Gy). Data is shown with mean ± SEM from 3 independent experiments. *p<0.1 and ****p < 0.0001 (one-way ANOVA, followed by Kruskal-Wallis test). F. Levels of RPA70 and phospho-RPA32 (pS4+pS8) in RPE1 hTERT *TP53^−/−^* cells of the genotypes indicated treated with or without 10 Gy ionising radiation (IR). Antibodies against vinculin were used as a loading control. G. Quantification of the number of phospho-RPA32 (pS4+pS8) foci in S-phase RPE1 hTERT *TP53^−/−^* cells with the *BRCA1* and *SCAF1* genotypes indicated, determined by PCNA staining before and after exposure of cells to ionising radiation (IR; 10 Gy). Data is shown with mean ± SEM from 3 independent experiments. ns: p>0.9999 (One-way ANOVA, followed by Kruskal-Wallis test). UT, untreated.

**Table S1. Mass spectrometry summary statistics.** Numbers of identified and quantified peptide to spectrum matches (PSM), (phospho)peptides in the phosphopeptide-enriched samples from U2 O-S cells ± berzosertib or gartisertib.

**Table S2. Phosphoproteomic dataset: U2 O-S cells, HU ± berzosertib** Quantitative proteomics dataset showing all phospho–peptides identified in U2 O-S cells treated with HU ± berzosertib. Phospho–peptides with identical sequence and post–translational modifications which were detected in the same chromatography fraction, had their respective TMT channel intensities averaged and given unique identifier numbers (column A “peptide number”). A given peptide with a unique identifier number can appear multiple times in the table if there is more than a single phosphorylation site and/or MaxQuant assigns a range of possible phosphorylation site localisation probabilities for a site (values listed under “PTM score probability”, column I). For statistical analysis, each peptide with a unique identifier number was considered exactly once, independent of the number of entries in this table. A phosphorylation localisation site probability threshold of 0.994 was applied to give an FLR cut-off of 1% (11). The mass spectrometry nuclear phospho-proteomics data have been deposited to the ProteomeXchange Consortium via jPOSTrepo (PXD040469).

**Table S3. Phosphoproteomic dataset: U2 O-S cells, HU ± gartisertib** Quantitative proteomics dataset of all showing all phospho–peptides identified in U2 O-S cells treated with HU ± gartisertib. Phospho–peptides with identical sequence and post–translational modifications which were detected in the same chromatography fraction, had their respective TMT channel intensities averaged and given unique identifier numbers (column A “peptide number”). A given peptide with a unique identifier number can appear multiple times in the table if there is more than a single phosphorylation site and/or MaxQuant assigns a range of possible phosphorylation site localisation probabilities for a site (values listed under “PTM score probability”, column I). For statistical analysis, each peptide with a unique identifier number was considered exactly once, independent of the number of entries in this table. A phosphorylation localisation site probability threshold of 0.994 was applied to give an FLR cut-off of 1% (11). The mass spectrometry nuclear phospho-proteomics data have been deposited to the ProteomeXchange Consortium via jPOSTrepo (PXD040469).

**Table S4.** Overlap between hits in Table S2 (HU ± berzosertib) and Table S3 (HU ± gartisertib). Overlap is based on phosphorylation sites less abundant under (HU ± inhibitor) and independent of the actual detected peptide. Phosphorylated amino acid is shown surrounded by square brackets in column K (“Motif”).

**Table S5.** New ATR targets already implicated in cell responses to DNA damage or replication stress. Listed are phospho-sites which less abundant common to both (HU ± inhibitor) experiments. In case phospho-site appears in different peptides, these are marked in grey. Phosphorylated amino acid is indicated in red in column K (“Motif”).

**Table S6.** New ATR targets not previously implicated in cell responses to DNA damage or replication stress. Listed are phospho-sites which less abundant common to both (HU ± inhibitor) experiments. In case phospho-site appears in different peptides, these are marked in grey. The phosphorylated amino acid is preceded by a “p” in column F (“Peptide sequence”).

**Table S7.** Lists of the reagents, antibodies, plasmid constructs, siRNA sequences, peptide sequences and primer sequences used in this study. Datasheets for each plasmid used in this study will be available on a dedicated page of our Reagents website upon publication.

